# MethylationToActivity: a deep-learning framework that reveals promoter activity landscapes from DNA methylomes in individual tumors

**DOI:** 10.1101/2020.06.09.143172

**Authors:** Justin Williams, Beisi Xu, Daniel Putnam, Andrew Thrasher, Chunliang Li, Jun Yang, Xiang Chen

**Affiliations:** Department of Computational Biology, St. Jude Children’s Research Hospital, Memphis, Tennessee, United States; Center for Applied Bioinformatics, St. Jude Children’s Research Hospital, Memphis, Tennessee, United States; Department of Tumor Cell Biology, St. Jude Children’s Research Hospital, Memphis, Tennessee, United States; Department of Surgery, St. Jude Children’s Research Hospital, Memphis, Tennessee, United States

**Author notes:** Corresponding author: Xiang Chen Department of Computational Biology St. Jude Children’s Research Hospital 262 Danny Thomas Place Mail Stop 1135 Memphis, TN 38105 U.S.A.

**Keywords:** DNA methylation, histone modifications, convolutional neural network, transfer learning

## Abstract

Although genome-wide DNA methylomes have demonstrated their clinical value as reliable biomarkers for tumor detection, subtyping, and classification, their direct biological impacts at the individual gene level remain elusive. Here we present MethylationToActivity (M2A), a machine learning framework that uses convolutional neural networks to infer promoter activities (H3K4me3 and H3K27ac enrichment) from DNA methylation patterns for individual genes. Using publicly available datasets in real-world test scenarios, we demonstrate that M2A is highly accurate and robust in revealing promoter activity landscapes in various pediatric and adult cancers, including both solid and hematologic malignant neoplasms.

## Background

Transcriptional regulation is fundamental to the identity and function of cells. Deregulation of gene expression is a defining feature of common diseases, including cancers. Promoters, the regulatory regions surrounding the transcription starting sites (TSSs), integrate signals from distal enhancers and local histone modifications (HMs) to initiate transcription. Almost half of human protein-coding genes harbor multiple TSSs; consequently, promoter activities determine both the level of transcription and the transcript isoforms that are expressed, with the latter potentially having different translation efficiencies and encoding different protein sequences [1]. Tumors frequently use alternative promoters to increase the isoform diversity [2, 3], to activate oncogenes that are normally repressed [1–3], and to evade host immune attacks by immunoediting [3, 4]. Compared to cancers in adults, pediatric tumors harbor fewer mutations [5, 6] and use epigenetic deregulation to promote tumorigenesis and progression [7].

Promoter activities can be determined experimentally through transcriptomic approaches, such as CAP analysis gene expression (CAGE), or through epigenomic approaches, including chromatin immunoprecipitation followed by sequencing (ChIP-seq) [8]. Because of transcript degradation by 5′ RNA exonucleases, ChIP-seq approaches for specific HMs have been the gold standard for studying promoter activities [9]. Several studies [10–12] have demonstrated that HMs and other epigenetic features can be used to predict gene expression. Using a linear regression model, Karlić et al. show that approximately 50%–60% of the variation in gene expression can be accounted for, and that ∼50% of the variation in gene expression can be modeled by promoter H3K27ac enrichment alone [10]. Subsequent work by Dong et al. further explained 69% of gene expression variance using a hybrid random forest/linear regression model with features derived from 11 HMs, one histone variant, and DNase I hypersensitivity [11]. More recently, Singh et al. used deep-learning models on five HMs to predict gene expression status (high/low) and achieved an average AUC of 0.80 [12]. However, the scarcity of pediatric tumors, the limited amounts of fresh starting material available, and the extensive workload involved in acquiring the promoter activity landscapes constrain their interrogation for individual patient tumors [13, 14].

DNA methylation (DNAm) is a well-studied, relatively stable, and inheritable epigenetic regulatory mechanism that involves transferring a methyl group to cytosine (C) to form 5-methylcytosine (5mC), mostly in the CpG context. In contrast to HMs, DNAm can be accurately and robustly profiled in various tissues, including archival formalin-fixed, paraffin-embedded (FFPE) tumor samples, through both array [15, 16] and sequencing [17] platforms; therefore, it has exceptional applicability to studying epigenetic deregulation in tumors. Consequently, genome-wide DNAm profiles represent a widely available epigenetic asset for studying epigenetic abnormalities in primary tumors.

The DNAm pattern is mechanistically connected with transcription factor binding and HMs [18–25]. It also plays critical roles in establishing the chromatin structure in physiologic and pathologic conditions [26, 27]. Moreover, recent applications of machine learning to genome-wide DNAm patterns have demonstrated that DNAm can accurately predict the patterns of chromatin packaging (A/B compartments, the square of the Pearson correlation coefficient *R*^2^ = 0.50–0.66) [28–30] and can reveal distinct subgroups with prognostic significance among patients with cancer [31, 32]. Recently, DNAm signature–based molecular classifiers were shown to improve diagnostic accuracy, as compared to that of traditional schemes, further demonstrating the critical regulatory roles of DNAm in tumor development [33, 34]. However, unlike HMs, where established biological interpretations of various marks have resulted in a general “histone code” hypothesis [35, 36], the relation between DNAm signatures and their transcriptional regulatory roles is complex and nonlinear. In many cases, even promoter DNAm may positively and negatively correlate with gene expression depending on the genomic structure involved in a given tumor [37]. Consequently, with few exceptions (e.g., hypermethylation of the promoters of *RB1*, *CDKN2A*, and *MGMT*) [38], the contribution of DNAm to the regulation of expression of individual genes remains largely elusive [39–41]. Recent attempts to use DNAm signatures to account for gene expression levels have had limited success, with the best model (binomial distribution probit regression [BPR] model) capturing 25%–49% of the expression variations [42]. Undoubtedly, the lack of interpretability of the DNAm pattern at the individual gene level has severely hampered our understanding of the biological significance of DNAm signatures.

To address these challenges, we have developed MethylationToActivity (M2A), a deep-learning framework. The central hypothesis of M2A is that the complex relation between DNAm signatures and promoter activities (measured as H3K4me3 and H3K27ac enrichment in the TSS ± 1 kb region) can be captured by incorporating both summary statistics extracted from window-based CpG methylation levels and high-order spatial information from these windows in the promoter and flanking regions (up to 25 kb from the TSS). Using a cohort of six pediatric neuroblastoma (NBL) orthoptic patient-derived xenograft (O-PDX) samples profiled in the Pediatric Cancer Genome Project (PCGP) [43], we trained the model using whole-genome bisulfite sequencing (WGBS) data to predict the enrichment of H3K4me3 and H3K27ac (two HMs critical for promoter [10]) for genome-wide annotated promoters. We validated the predictive accuracy of the model in the remaining NBL samples (N = 10, WGBS) from the same cohort. We further confirmed its accuracy and generalizability in diverse tumor types from four publicly available datasets representing real-world applications, including (1) pediatric rhabdomyosarcoma (RMS) O-PDX tumors profiled in the Pediatric Cancer Genome Project (N = 16, WGBS) [44]; (2) a set of commonly used cell lines profiled in ENCODE (N = 9, WGBS) [45]; (3) primary acute myeloid leukemias (AMLs) profiled by the BLUEPRINT consortium (N = 19, WGBS) [46]; and (4) a large primary Ewing sarcoma (EWS) cohort using reduced representation bisulfite sequencing (RRBS) (N = 140) [22]. These applications demonstrate that M2A can accurately reveal promoter activities from DNAm patterns, which will be of great use not only in functionally interpreting differential DNAm patterns but also in profiling promoter usage in individual patient tumors. This will facilitate precision medicine by tailoring treatments based on both genetic variants and epigenetic deregulations.

## Results

### Extensive diversity of promoter activity among *MYCN-*amplified NBL cell lines and O-PDX models

To date, most cancer HM profiling studies have made use of tumor models, including cell lines, xenografts, and more recently, organoids. Technical limitations and challenges when working with human tumor tissues prevent the generation of high-quality ChIP-seq profiles for primary patient specimens [47]. Despite the documented epigenetic heterogeneity [48], a common practice in deciphering major HM deregulations in various cancers is to extrapolate the epigenetic profiles from related cancer models (surrogate models). Many studies [43, 44, 49–52] have compared model systems to primary tumors with respect to characteristics such as mutations, gene expression, and DNAm signatures. In this study, we began by evaluating the level of promoter activity diversity in closely related NBL models. Specifically, we evaluated promoter activity, as measured by the H3K27ac level, in three O-PDX models (SJNBL046, SJNBL108, and SJNBL013763) and three cell line models (IMR-32, NB-5, and SKNBE2) that harbor *MYCN* amplification with no other major oncogenic mutations. All samples displayed a bimodal distribution of promoter H3K27ac levels across the genome (Additional file 1: Figure S1), and O-PDX models had a marginally higher fraction of active promoters (mean: 31.9%, range: 27.6%–36.1%) than did cell line models (mean: 26.1%, range: 25.6%–26.7%) (*P* = 0.14, Student’s *t*-test). However, there were extensive variations in the promoter activities in both the cell line models (Figures 1a and 1d) and the O-PDX models (Figures 1b and 1e). Moreover, greater divergence was observed between a cell line model and an O-PDX model (mean: 34.9%, range: 29.9%–39.0%) than between two cell line models (mean: 31.0%, range: 29.2%–32.1%; *P* = 0.44, Student’s *t*-test) or between two O-PDX models (mean: 31.0%, range: 22.9%–37.0%; *P* = 0.02, Student’s *t*-test) (Figure 1f). Variations in promoter activity may play a significant role in the transcriptional deregulation of individual tumors, as a substantial fraction of established cancer consensus genes (22.4% in O-PDX models and 31.1% in cell line models, including *APOBEC3B*, *TGFBR2*, *PAX7*, *HOXA11*, *PDCD1LG2*, *PTK6*, *BCL11B*, *FAS*, and *MYC*; (Additional file 2: Table S1) displayed heterogeneous promoter activities in the surveyed tumor models. Therefore, we sought to develop a computational approach to infer the promoter activity landscape for individual tumors.

**Figure 1.**
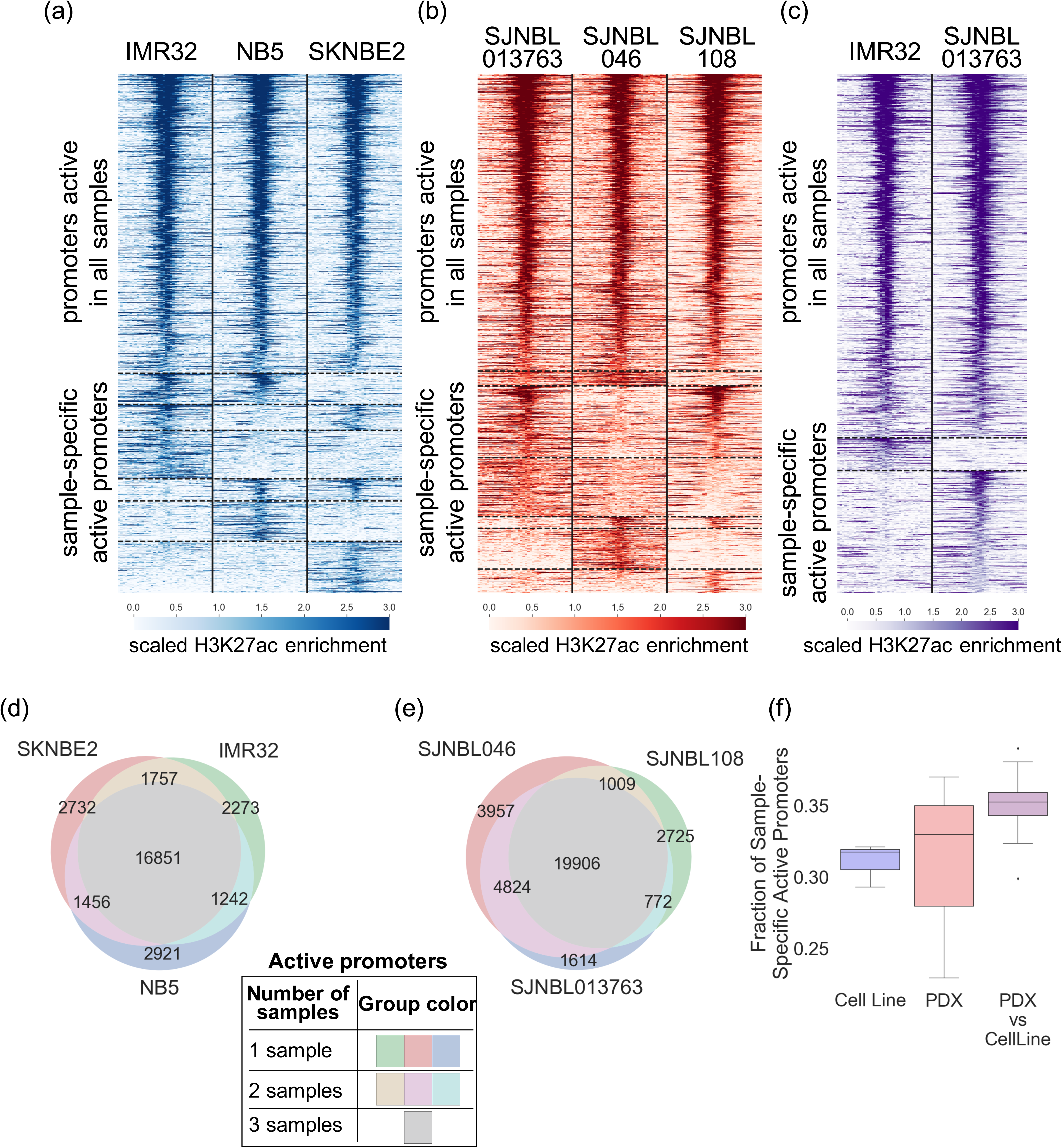
Variations of promoter H3K27ac levels among *MYCN* amplified NBL samples. (a−c) Promoters were classified as active or inactive based on H3K27ac levels in individual NBL tumors: **(a)** NBL cell line samples, **(b)** NBL O-PDX samples, and **(c)** NBL cell line and O-PDX samples. Each promoter region plotted spans TSS ± 5,000 bp binned by non-overlapping 250 bp windows; the color bar represents the scaled windowed H3K27ac enrichment, from 0 (lowest) to 3 (highest). Horizontal dotted lines delineate shared promoters (active in all samples) and sample-specific promoters (promoter activity in at least one sample was different from the remaining samples). Promoters were sorted by average descending H3K27ac enrichment across all samples within each group. **(d, e)** Venn diagram indicating the number of shared and sample-specific H3K27ac promoter activities between NBL cell line samples (**d**) and O-PDX samples **(e)**. **(f)** The promoter activity variation is highlighted by the proportion of sample-specific active promoters among all H3K27ac active promoters, as compared within cell line or O-PDX NBL samples, and between cell line and O-PDX NBL samples.

### M2A: a deep-learning framework to reveal promoter activities from DNA methylation

DNAm plays a critical role in determining the framework of gene expression for a given cell/cellular state. However, the highly complex and non-linear relations between DNAm patterns and HMs severely hamper the interpretability of the biological impact of differential DNAm patterns. Previous studies have shown the usefulness of extracting higher-order methylation features [42], for predicting gene expression. Moreover, recent studies applied deep-learning approaches to infer DNAm states from their local sequence composition and adjacent DNAm states [53]. We hypothesize that these high-level DNAm features (that capture the spatial information from DNAm patterns in the promoter and regions in its vicinity) could also provide an opportunity to infer promoter activities such as H3K27ac and H3K4me3 enrichment accurately. We propose to use a convolutional neural network (CNN)–based deep-learning framework to extract such features.

The M2A conceptual framework and workflow is shown in Figure 2. M2A starts with raw DNAm feature extraction from around individual TSSs (Figure 2a). This is followed by high-level feature extraction through the CNN layers and mapping between the generalized feature and the final output (i.e., the H3K4me3 and H3K27ac of the promoter) in the fully connected (FC) layers. The vanilla model described in this report was trained on six NBL PDX tumors (SJNBL046_X, SJNBL013761_X1, SJNBL012401_X1, SJNBL013762_X1, SJNBL013763_X1, and SJNBL015724_X1; Figure 2b) for which comprehensive genomic and epigenomic profiling data are available, including the results of whole-genome sequencing, whole-exome sequencing, RNA sequencing, WGBS, and ChIP-seq of eight histone marks (H3K4me1, H3K4me2, H3K4me3, H3K27me3, H3K27ac, H3K36me3, H3K9/14ac, and H3K9me3), CTCF, BRD4, and RNA polymerase II (PolII).

**Figure 2.**
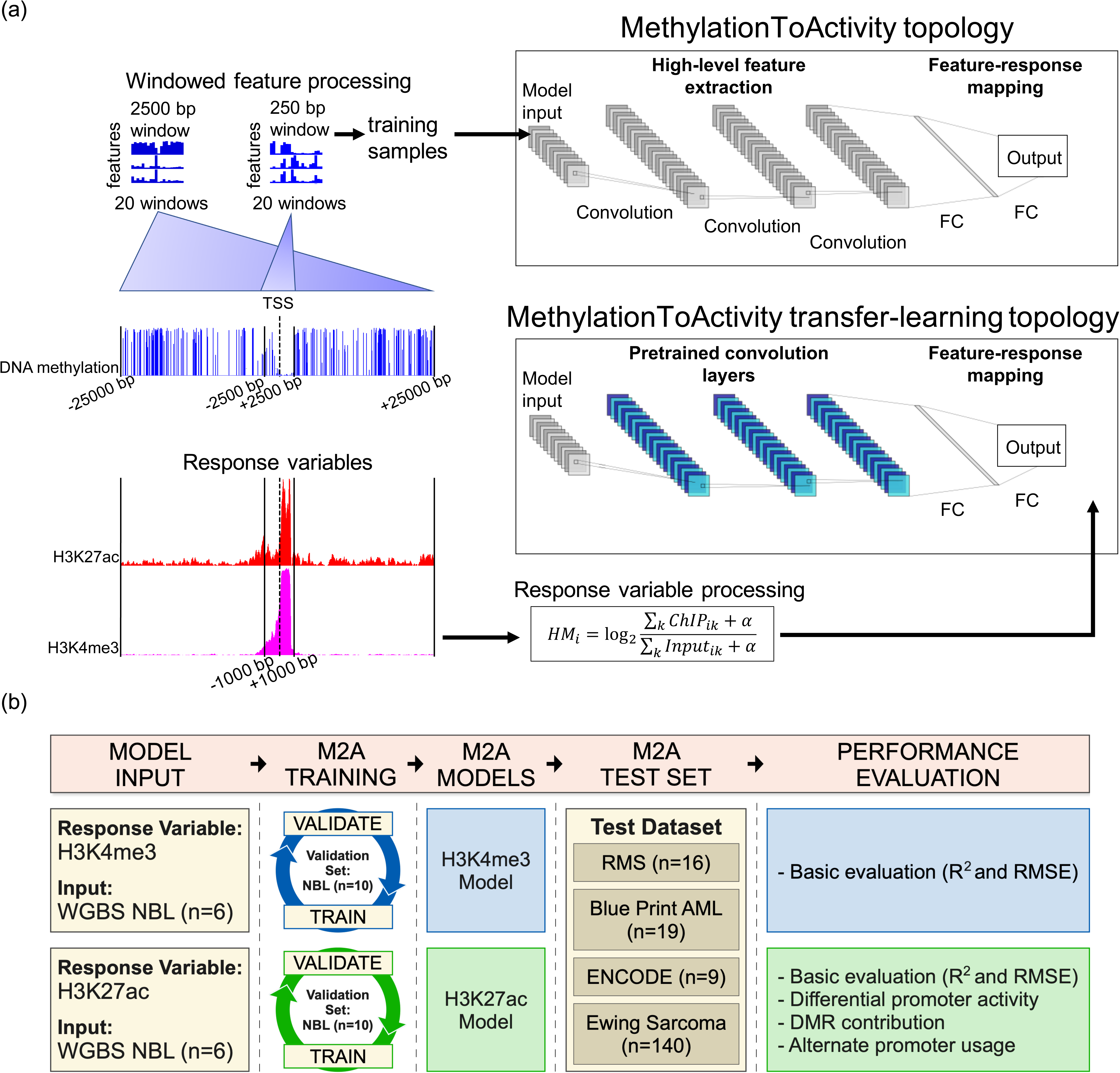
M2A feature processing and training workflow. The M2A framework hinges on the feature processing pipeline. **(a)** First, windowed features (20 total non-overlapping windows for each of two sizes including 250 bp and 2,500 bp) centered around the TSS are calculated from WGBS data for each unique promoter region, extending up to 2,500 bp (250 bp window), and 25 kbp (2,500 bp window) away from the TSS. Response variables (H3K27ac and H3K4me3) for separate model training were generated for matching promoter regions (TSS ± 1 kbp). The matching window features and response variables serve as input to the model topologies, where M2A first extracts high-level features by using a series of convolutional layers then maps these features to response variables in fully-connected (FC) layers. Transfer learning with M2A leverages pretrained feature extraction (frozen CNN layers indicated in blue), training only the FC layers. **(b)** The overall workflow for training, validation, and testing M2A is detailed, as well as an overview of the analyses performed to validate M2A performance in different real-world applications. M2A models for H3K4me3 and H3K27ac were trained separately, indicated by blue (H3K4me3) and green (H3K27ac).

We started with an analysis of the information content in DNAm patterns by examining the input feature distribution in different windows, among active (high H3K27ac), poised (high H3K4me3 and low H3K27ac), and inactive promoters (low H3K4me3 and low H3K27ac) in the six NBL O-PDX training samples. These features show distinct patterns among the three promoter categories (Additional File 1: Figure S2), indicating the feasibility of modeling promoter activities from DNAm patterns. Although the interpretability of CNN extracted features remains an active field of research in deep-learning [54], we examined the efficacy of CNN extracted features in modeling the promoter activities. We first compared the square of Pearson’s correlation (*R^2^*) between each feature (both raw input and CNN extracted features) and the response variable (H3K27ac) in the training set, The analysis revealed that CNN-extracted features have significantly higher *R^2^* with the response (250 bp: *P* = 1.5 × 10^−11^, 2500 bp *P* = 3.9 × 10^−5^, Wilcoxon signed-rank test, Additional File 1: Figure S3). We further evaluated the best features for both raw input and CNN-extracted features in the validation samples and again the CNN-extracted features significantly outperformed the raw input features (250 bp: *P* = 1.1 × 10^−5^, 2500 bp *P* = 1.1 × 10^−5^, Wilcoxon signed-rank test, Additional File 1: Figure S3).

### M2A produces a highly accurate landscape of promoter activity in pediatric NBL

To evaluate the performance of M2A, we first explored its performance in the remaining NBL samples in the cohort (the validation set), including one O-PDX tumor, one primary autopsy tumor, and eight cell lines. Using the validation set, we compared the performance of the M2A framework of three CNN layers and two FC layers (Figure 2) with three frequently used statistical and machine learning approaches (baseline models), namely multivariate adaptive regression splines (MARS), random forest, and artificial neural network (ANN) consisting of only two FC layers. In every instance, the M2A framework outperformed baseline models (Additional file 2: Table S2). From a qualitative perspective, M2A correctly revealed the bimodal distribution of the promoter activities for both H3K4me3 and H3K27ac in all samples, and from a quantitative perspective, the inferred genome-wide promoter activity landscape was highly accurate for individual samples for both H3K4me3 (*R*^2^ = 0.933 ± 0.019; RMSE = 0.621 ± 0.072) (Figures 3a and 3d) and H3K27ac (*R*^2^ = 0.799 ± 0.053; RMSE = 0.644 ± 0.074) (Figures 3b, and 3c). Moreover, the addition of CNN layers was merited, as there was a decrease in the prediction error (measured as 1 − *R*^2^) from the next highest performer by 17.8% (*P* = 0.0020, Wilcoxon signed-rank test) and 12.4% (*P* = 0.0020, Wilcoxon signed-rank test) for the model topologies for H3K4me3 and H3K27ac, respectively (Additional file 2: Table S2).

**Figure 3.**
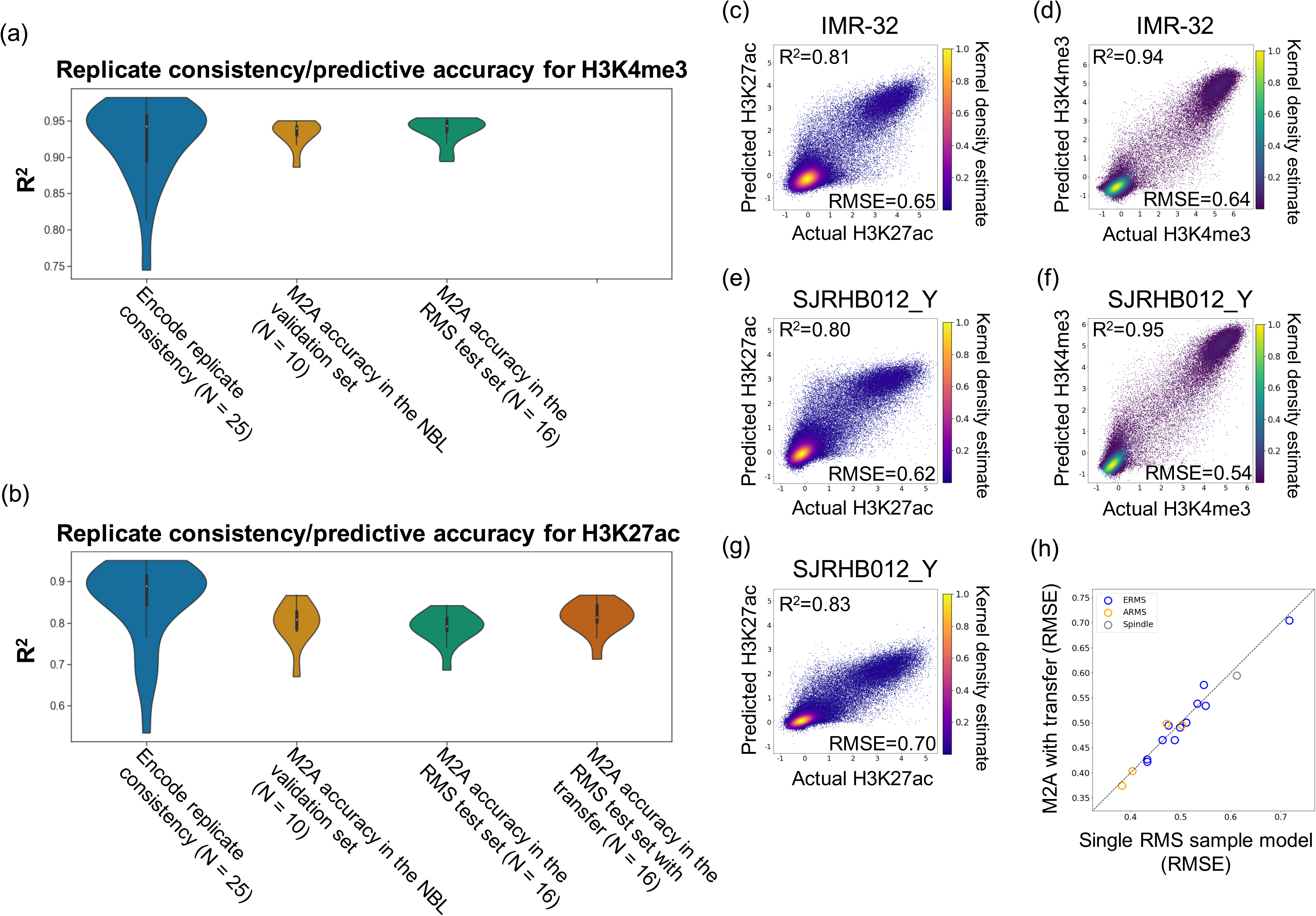
M2A performance in NBL and RMS cohorts. (a, b) Analysis of the performance of M2A in NBL and RMS cohorts with **(a)** H3K4me3 inference and **(b)** H3K27ac inference. ENCODE replicate consistencies were calculated as Pearson’s correlation squared (*R^2^*) between replicates (two replicates per sample). ENCODE sample KMS-11 was excluded as an apparent outlier *R^2^*=0.016, RMSE = 1.869. Prediction accuracy was measured by *R^2^* between the actual measurement and M2A’s prediction. **(c−g)** Individual examples of median M2A performers in **(c)** NBL cell line H3K27ac inference, **(d)** NBL cell line H3K4me3 inference, **(e)** RMS H3K27ac inference (pre-transfer), **(f)** RMS H3K4me3 inference, and **(g)** RMS H3K27ac inference (post-transfer). To indicate the density of data points where individual data points cannot be resolved, a KDE was applied, called from 1 (highest) to 0 (lowest). **(h)** The boost to M2A performance (measured by RMSE) due to transfer learning is shown, as applied and tested in RMS samples.

Our analysis of *MYCN*-amplified NBL cell line and O-PDX models has revealed substantial variations in their promoter activities, which is a potential caveat to the practice of surrogate model (representing primary tumor epigenomes by a few profiled models). Conversely, M2A produced highly accurate promoter activity landscapes, significantly outperforming the observed consistency between training and testing samples for both H3K4me3 (*R*^2^ = 0.891 ± 0.023, *P* = 2.3 × 10^−5^, Wilcoxon rank-sum test) and H3K27ac (*R*^2^ = 0.720 ± 0.045, *P* = 9.5 × 10^−5^, Wilcoxon rank-sum test; Additional file 2: Table S3). Remarkably, in nine (of 10) test samples, the accuracy of the M2A-inferred promoter H3K27ac activity was better than the highest similarity attained by any individual training sample (*P* = 0.027, Wilcoxon signed-rank test). The same pattern was observed for H3K4me3 levels, with M2A being more accurate for nine of 10 samples (*P* = 0.037, Wilcoxon signed-rank test), demonstrating the accuracy of M2A in revealing individual tumor promoter activity landscapes. Finally, the predictive accuracy of M2A was comparable to the experimental consistency observed between replicates from the same cell lines profiled in ENCODE for H3K4me3 (*R*^2^ = 0.933 ± 0.018 for M2A vs. 0.922 ± 0.056 for ENCODE replicates [N = 25]; *P* = 0.55, Wilcoxon rank-sum test) (Figure 3a; Additional file 2: Table S4). The accuracy of M2A also approached the replicate consistency for H3K27ac (*R*^2^ = 0.799 ± 0.050 for M2A vs. 0.849 ± 0.047 for ENCODE replicates [N = 26]; *P* = 0.0078, Wilcoxon rank-sum test) (Figure 3b: Additional file 2: Table S4). Measurement of the root mean square error (RMSE) revealed a similar pattern (Additional file 2: Table S4).

### M2A is generalizable and scalable

Aside from the model accuracy, there are two additional requirements with practical importance for deploying a machine learning model (such as M2A) in real-world applications: (1) generalizability, i.e., M2A needs to achieve a similar performance with a set of unseen test samples, including tumor/tissue types not used in the model training; and (2) scalability, i.e., M2A must be able to be applied efficiently to external data.

We first demonstrated the accuracy, generalizability, and scalability of M2A by using test samples from rhabdomyosarcoma (RMS) O-PDX tumors. The RMS O-PDX dataset consists of 16 pediatric RMS tumors (11 embryonal, four alveolar, and one spindle subtype, termed ERMS, ARMS, and spindle subtypes, respectively). As with the NBL cohort, each RMS sample was extensively profiled, including by WGBS, RNA-seq, and ChIP-seq of H3K4me3 and H3K27ac. Using the vanilla M2A model (the 3CNN-FC model trained on the six NBL PDX samples), M2A achieved an overall predictive accuracy with the RMS dataset that was comparable to that of the NBL test group for both H3K4me3 (*R*^2^ = 0.937 ± 0.017, *P* = 0.30; RMSE = 0.639 ± 0.119, *P* = 0.82, Wilcoxon rank-sum test) (Figures 3a and 3f; Additional file 2: Table S5) and H3K27ac (*R*^2^ = 0.790 ± 0.037, *P* = 0.44; RMSE = 0.589 ± 0.084, *P* = 0.058, Wilcoxon rank-sum test) (Figures 3b and 3e; Additional file 2: Table S5), which was comparable to or significantly outperformed the observed similarities between two different RMS tumors for H3K4me3 (*R*^2^ = 0.917 ± 0.028, *P* = 0.0020; RMSE = 0.646 ± 0.133, *P* = 0.64, Wilcoxon rank-sum test) and H3K27ac (*R*^2^ = 0.780 ± 0.066, *P* = 0.43; RMSE = 0.550 ± 0.095, *P* = 0.14, Wilcoxon rank-sum test). The accuracy of the inferred H3K4me3 activity was comparable to the inter-replicate consistency of the ENCODE samples (*P* = 0.83, Wilcoxon rank-sum test).

By definition, generalizability can be achieved only in the absence of over-fitting (or “memorization” of the training data). Neural networks often fall victim to this problem through a combination of factors, including relatively small training datasets and/or over-parameterization. The relatively consistent expression of housekeeping genes across different tissues may lead to an inaccurate (often inflated) interpretation of the performance measurement in such a model, as evidenced by the relatively high *R*^2^ value (0.663 ± 0.040) between the promoter H3K27ac level of a random RMS test tumor and the most similar NBL training tumor (Additional file 1: Figure S4a). Therefore, we focused on the set of genes that are differentially expressed (DE) in RMS and NBL PDX samples [51], for which an over-fitted or memorized model would perform poorly. Not surprisingly, the average correlative consistency between the NBL validation samples and the most similar NBL training sample dropped from 0.755 to 0.599 when the measurement was restricted to promoters encoding the DE genes (Additional file 1: Figure S4b), whereas a sharp decline (from 0.663 to 0.259) was also observed for RMS test tumors (Additional file 1: Figure S4b). Conversely, the six-PDX NBL-trained M2A model maintained high accuracy for promoters of DE genes in both the NBL validation set (*R*^2^ = 0.729 ± 0.071 and the RMS test set (*R*^2^ = 0.715 ± 0.044) (Additional file 1: Figure S4b), further demonstrating the generalizability of M2A.

M2A is efficient and scalable. For a local implementation of M2A (source code, built models and a Docker image available at https://github.com/chenlab-sj/M2A), the training of the vanilla M2A model (with six NBL O-PDX tumors) takes approximately 16 min (using a Tesla P100-16GB GPU). The feature extraction and promoter activity prediction from WGBS data (as a genome-wide DNAm level file in a tab-delimited text format) can be executed on a personal computer (in this case, we used a MacBook Air with a 2.2-GHz Intel Core i7 and 8-GB 1600-MHz DDR3 RAM) and takes 15–19 min. Moreover, we have implemented a cloud version of M2A (https://platform.stjude.cloud/workflows/methylation-to-activity), available to the general research community.

### Transfer learning further improves the performance of M2A with minimal additional input in the target domain

Although we have demonstrated the generalizability of M2A in the RMS dataset, the fact that epigenetic genes are frequently mutated in pediatric tumors [7] raises the possibility that individual tumor types carry a type-specific interpretation of the DNAm patterns. When ChIP-seq measurement is available for sufficient samples, a type-specific model is desirable. However, although pediatric solid tumors as a group constitute a rare disease, they comprise many different tumor types, and it is rare to have sufficiently profiled samples available for many of them. In addressing this challenge, we hypothesize that a fixed feature-extraction strategy (transfer learning) can achieve the goal of deriving an efficient tumor type–specific model by using a small labeled dataset. A primary assumption here is that generalized features extracted based on a large dataset are similarly informative for apparently different tasks. The feature learning and selection characteristics of CNNs provide exceptional portability in various tasks with extremely small labeled datasets.

In M2A, the CNN layers capture generalized DNAm features and the FC layers learn the mapping function between the DNAm features and the promoter activities. Here we start with the pretrained vanilla M2A model, fix the feature-extraction layers (CNN layers), and use a single sample from the target tumor type to update the mapping function (the weights and biases of the FC layers). Because the consistency of M2A for H3K4me3 approached the inter-replicate consistency in both NBL and RMS datasets, we focused on H3K27ac inference for transfer learning. Upon performing transfer learning with a single sample in the RMS dataset, we observed significantly improved accuracy (*R*^2^ = 0.813 ± 0.038, *P* = 3.1 × 10^−5^, Wilcoxon signed-rank test) (Figures 3b, 3g, and 3h; Additional file 2: Table S5). Moreover, this model significantly outperformed a single RMS sample model with the identical model architecture, in which both the CNN layer and the FC layers were derived from the RMS training sample (*P* = 9.2 × 10^−5^, Wilcoxon signed-rank test) (Additional file 1: Figure S5; Additional file 2: Table S6) and marginally outperformed the observed similarities between different RMS tumors (*P* = 0.053, Wilcoxon rank-sum test). This analysis demonstrated the value of both the pretrained CNN layers for general feature extraction and a single profiled sample in the target domain. Consequently, we applied transfer learning to both the EWS and AML datasets. However, transfer was not feasible in the ENCODE dataset because those cell lines were derived from different tissues.

### M2A accurately reveals promoter activity landscapes in adult tumors and in hematologic malignant neoplasms

We next evaluated the performance of M2A in independently collected datasets, including ones for adult tumors and hematologic malignant neoplasms. Upon analyzing nine ENCODE cell lines, we found that differences in antibody usage and experimental protocols between the ENCODE and NBL datasets resulted in different signal-to-noise profiles (Additional file 1: Figure S6), with higher RMSE values between the model predictions and the actual observations (H3K4me3: 0.961 ± 0.238, *P* = .0019; H3K27ac: 0.918 ± 0.188, *P* = 2.6 × 10^−4^, Wilcoxon rank-sum test). However, the predicted activities remained highly correlated with the experimental measurements (H3K4me3: *R*^2^ = 0.895 ± 0.027; H3K27ac: *R*^2^ = 0.680 ± 0.149) (Additional file 1: Figure S7). Although the accuracy of H3K4me3 was relatively uniform, H1-ESC and SK-N-SH were outliers with a substantially lower accuracy of H3K27ac inference (by the boxplot-based method [55]) (Additional file 2: Table S4; Additional file 1; Figure S8a). An investigation of the promoter activity (the measured and inferred H3K27ac levels) and the measured gene expression in H1-ESC revealed that a subset of actively transcribed genes showed little or no H3K27ac levels in their promoters, where M2A inferred relatively strong promoter activity (Additional file 1: Figure S8b). Consequently, the inference of promoter activity by M2A outperformed the actual measurement in terms of both the quantitative consistency with the gene expression level (*R*^2^ = 0.536 for the M2A-inferred H3K27ac level vs. 0.435 for the measured H3K27ac level) (Additional file1: Figures S8b and S8c) and the accuracy in predicting expressed genes (AUC = 0.891 for the M2A-inferred H3K27ac level and 0.881 for observed H3K27ac level) (Additional file 1: Figure S9d). We also observed a small fraction of inferred active promoters without strong expression; these may represent genes subject to transcriptional pausing (where transcription is initiated but there is no elongation), a distinctive feature of undifferentiated stem cells [56]. Similarly, better consistency with gene expression levels was observed in SK-N-SH (Additional file 2: Table S7).

We further evaluated the performance of M2A in revealing promoter activities in 19 acute myeloid leukemia (AML) primary patient samples collected by the BLUEPRINT consortium, which is part of the International Human Epigenome Consortium (IHEC) [46]. Analyses of observed promoter activity (measured by H3K27ac level) and gene expression (measured in fragments per kilobase of transcript per million mapped reads [FPKM]) in the same sample revealed non-uniform qualities with a wide range of consistency (mean *R*^2^ = 0.539, range: 0.178–0.720) (Additional file 2: Table S8). Similarly, the M2A-inferred promoter activity landscape displayed substantial variability with respect to the observed activities among these samples (mean *R*^2^ = 0.473, range: 0.031–0.729). Strikingly, the consistency between the observed promoter activity and gene expression was highly predictive of the performance of M2A with individual samples (*R*^2^ = 0.975, *P* = 4.1 × 10^−15^, Pearson’s correlation test) (Additional file 1: Figure S10). Finally, although gene expression was not used in model generation with M2A (for the vanilla model or the transfer learning step), the promoter activity inferred by M2A showed uniform consistency with gene expression (mean *R*^2^ = 0.628, range: 0.541–0.684) (Additional file 2: Table S8).These results jointly suggest that the ChIP-seq library quality is a potential confounding factor for both the consistency between the observed (from ChIP-seq) and the M2A-inferred promoter activity landscapes and the consistency between the observed promoter activity and gene expression in these samples.

### Promoter activity landscape inferred by M2A faithfully recapitulates the subtype difference between embryonal and alveolar rhabdomyosarcomas

Identifying recurrent epigenetic deregulations (epi-drivers) is a primary research focus in cancer epigenome studies [57]. To this end, we investigated whether subtype-specific epigenetic deregulation was captured in M2A-revealed promoter landscapes in the RMS O-PDX tumors. A t-SNE embedding using M2A-inferred promoter activity landscapes (from the NBL-trained model) recapitulated the clear separation of ARMS and ERMS tumors (Figure 4a) in the DNAm profiles (data not shown), which further demonstrates the generalizability of the CNN-extracted high-order DNAm features. Importantly, when focusing on the promoters of DE genes in the ARMS and ERMS subtypes, the vanilla M2A model faithfully retained the subtype-specific promoter activity patterns (*R*^2^ = 0.713 for DE genes with a single annotated promoter; 0.621 when all annotated promoters for DE genes were included) (Additional file 1: Figures S11a and S11c). Transfer learning using data from a single RMS further improved the consistency (*R*^2^ = 0.758 and 0.673, respectively) (Additional file 1: Figures S11b and S11d).

**Figure 4.**
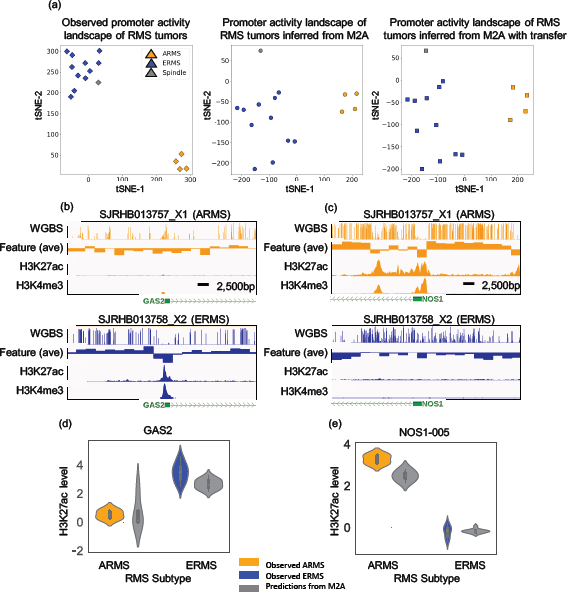
M2A recapitulates subtype differences in RMS. (a) A t-distributed Stochastic Neighbor Embedding (tSNE) analysis of observed (left), M2A inferred (center: pre-transfer, right: post-transfer) H3K27ac promoter levels. Embryonal and alveolar RMS subtypes are well separated in each analysis, demonstrating that M2A inferred H3K27ac levels maintain the delineation of RMS subtypes, consistent with the observed H3K27ac tSNE analysis. **(b–d)** The subtype-specific genes *GAS2* **(b)** and *NOS1* **(c)** show subtype distinct patterns of DNAm, H3K27ac, and H3K4me3 levels. The windowed average DNAm feature (2,500 bp windows, over the genomic region TSS ± 25 kb) is shown as example (partial) M2A input. These subtype differences were faithfully recapitulated by the M2A H3K27ac inferences for *GAS2* **(d)** and *NOS1* **(e)**.

*GAS2* is a gene selectively expressed in ERMS [44]. Although promoter hypomethylation was found in the ARMS tumors, the M2A model correctly predicted significantly stronger promoter activities in ERMS tumors (*P* = 0.01, Wilcoxon rank-sum test) (Figures 4b and 4d). Similarly, although both ERMS and ARMS tumors had *NOS1-005* promoter hypomethylation, strong promoter activity was predicted in ARMS tumors only (*P* = 0.0015, Wilcoxon rank-sum test) (Figures 4c and 4e), consistent with the ChIP-seq measurement.

### M2A reveals the contribution of differentially methylated regions to promoter activities

Although DNA methylation patterns, including differentially methylated regions (DMRs), are well-established biomarkers for diverse diseases and have revealed molecularly and clinically different subtypes for many cancers, their functional importance in individual gene regulation is less clear [39–41]. For example, many cancer-specific CpG island hypermethylation regions occur in genes that normally are not expressed or are expressed at only a low level [58]. Moreover, even in genes that are both differentially expressed and differentially methylated in different subtypes, up-regulated samples can be associated with hypomethylation, hypermethylation, or both [37, 44]. The observed nonlinear relation between DNA methylation and gene expression complicated the functional interpretation of specific DMRs. In this analysis, we interpret the functional roles of DMRs based on the promoter activities of their associated DE genes.

To summarize unambiguously the contribution of DMRs to differential promoter activities, we focused on 371 genes in ERMS and ARMS that have a single annotated promoter and that are both differentially expressed (197 are over-expressed in ERMS, 172 are over-expressed in ARMS) and differentially methylated (169 are hypomethylated, 128 are hypermethylated, and 74 have both hypomethylation and hypermethylation) in the two major RMS subtypes (Additional file 2: Tables S9 and S10) [44]. Among these genes, 140 promoters showed significantly higher H3K27ac measurements in the over-expressed subtype (FDR < 0.1, Wilcoxon rank-sum test), whereas 206 promoters had measurements that were significantly higher when the DNAm-based H3K27ac activity was measured. These 206 promoters included 118 of the 140 promoters identified using the observed signals. These results suggest that M2A can reveal the role of DMRs in modulating the promoter activities of affected genes in a context-specific manner.

### M2A identifies subtype-specific promoter usage encoding different protein isoforms in rhabdomyosarcoma

Alternative promoter usage is an important pretranslational mechanism for tissue-specific regulation as it affects the diversity of isoforms available. Recently, light was shed on the pervasiveness of alternative promoter usage in cancer; in some cases, promoter usage is a more accurate reflection of patient survival than is gene expression [2]. Among 10,835 active genes with multiple annotated promoters in the RMS dataset, we found 2,584 genes (24%) with alternative primary promoter usage among 16 samples. We focused on 562 genes that 1) were active in both the ERMS and ARMS subtypes and 2) had subtype-specific promoter usage. We explored the accuracy of M2A in predicting alternative promoter usage in ARMS and ERMS (Additional file 2: Table S11).

Based on measured promoter activities, 428 genes exhibited significant usage difference between the two subtypes (FDR < 0.1, Wilcoxon rank-sum test [used as the ground truth]). The M2A-inferred promoter activity landscape revealed 276 genes for which there was a significant difference in promoter usage between the subtypes (FDR < 0.1, Wilcoxon rank-sum test), and 210 of them matched the ground truth (precision = 0.76, recall = 0.49, F1 score = 0.60) (Additional file 2: Table S11).

PDZ Domain Containing Ring Finger 3 (*PDZRN3*) is a known target of the PAX3/7–FOXO1 fusion protein [59–61], which blocks terminal differentiation in myogenesis [62]. M2A predicted a subtype-specific promoter usage pattern in *PDZRN3*. Functional studies have shown that PDZRN3 regulates myoblast differentiation into myotubes through transcriptional and posttranslational regulation of Id2 [62]. Its over-expression in ARMS was primarily driven by the fusion protein binding adjacent to an alternative promoter (*PDZRN3-006*) located 191 kbp downstream of the canonical promoter (*PDZRN3-001*) (Figure 5a). The subtype-specific isoform usage is accompanied by DMRs of the alternative promoter and its immediate downstream regions and is further confirmed by RNA-seq read alignment (Figure 5a). Compared to the canonical isoform expressed in ERMS, the ARMS-preferred *PDZRN3-006* isoform lacks the RING-finger and Sina domains in the N-terminus and harbors a shorter PDZ domain. The isoform difference, as well as the differences in expression level, between subtypes may contribute to the impairment of myogenesis at different stages in the development of ARMS and ERMS tumors.

**Figure 5.**
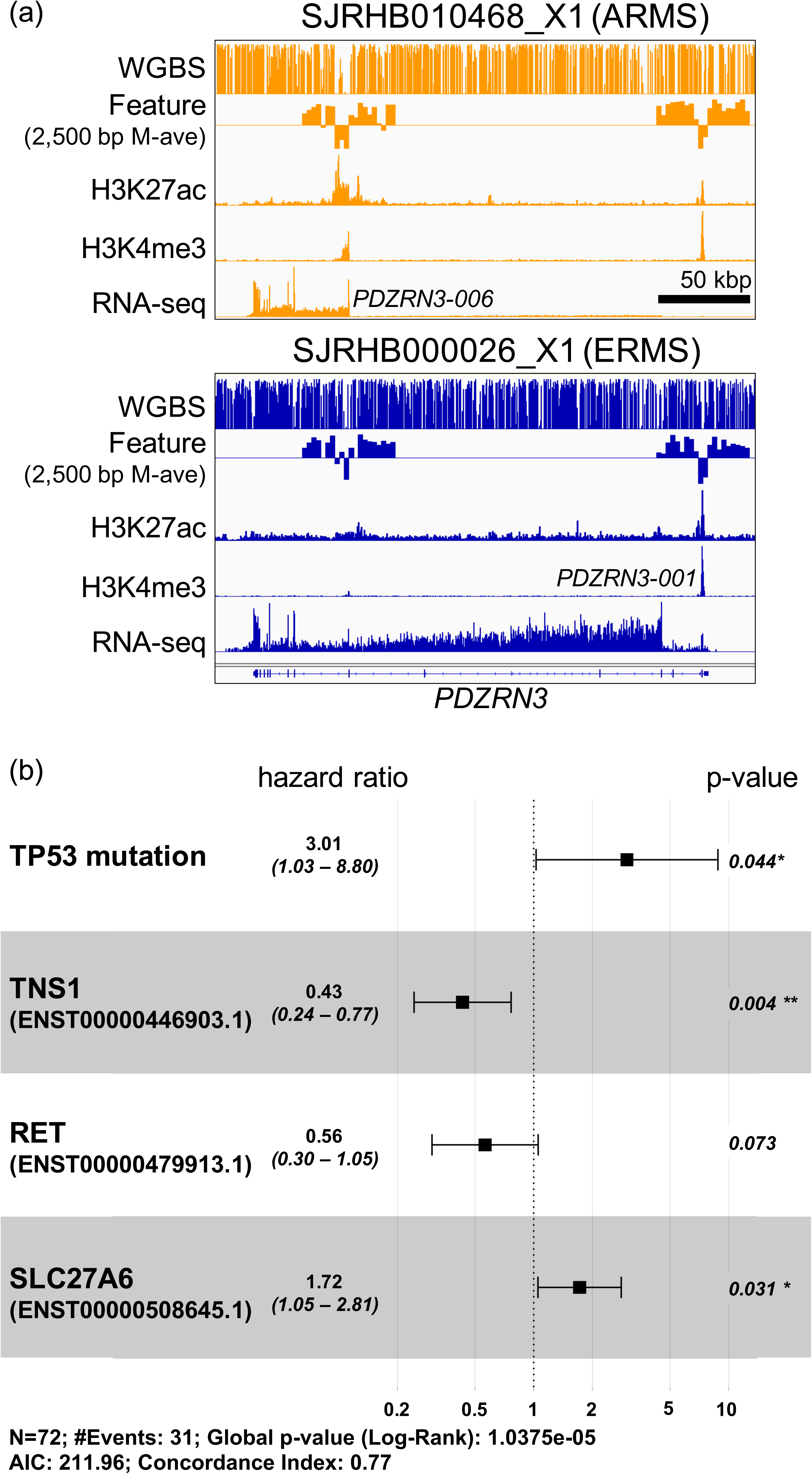
M2A reveals alternate promoter usage in RMS and EWS. (a) An analysis of alternate primary promoter usage between RMS subtypes ERMS and ARMS shows that M2A appropriately predicts subtype–specific promoter usage in *PDZRN3*, a known target of the PAX3/7–FOXO1 fusion protein, which is consistent at the observed values of H3K27ac, H3K4me3, RNA-seq, and DNA methylation. Partial M2A input (DNAm 2,500 bp windowed average) is shown to emphasize the DNAm patterns in the genomic region surrounding the TSS ± 25 kb. **(b)** Alternate promoter usage in EWS patient samples with and without *TP53* mutations were incorporated in a Cox proportional hazards model, highlighting the potential prognostic value of the isoforms identified by M2A.

### M2A identifies alternative promoter usages with potential prognostic values in Ewing sarcoma

We next examined the predicted epigenetic promoter activities in 140 EWS samples with DNAm data assayed by RRBS [22]. Three of these samples had matching ChIP-seq profiles for H3K4me3 and H3K27ac. To interrogate this dataset, we applied the pretrained vanilla model with transfer learning, as detailed above, to recalibrate the weights mapping the high-level features to the promoter activities in the EWS cohort. Despite the difference in DNAm platforms (WGBS for the NBL training model and RRBS for the EWS samples), the inferred promoter activity landscape was accurate (*R*^2^ = 0.718, 0.628, and 0.702 for the three samples with HM profiles, using leave-one-out prediction) (Additional file 2: Table S12).

Ewing sarcomas with mutations in *TP53* and *STAG2* have a particularly dismal prognosis [63]. We explored whether the promoter activity had additional prognostic value in 72 samples for which survival data was available. Because of the limited sample size, the initial analysis revealed a significant association between poor clinical outcomes and *TP53* mutations (*P =* 0.00047, log-rank test) (Additional file 1: Figure S12a) but not *STAG2* mutations (*P* = 0.67, log-rank test) (Additional file 1: Figure S12b). We identified 21 active genes that showed a potential difference (absolute mean difference of log-scaled activity≥1) between *TP53* mutant tumors and wild-type tumors, and we applied Cox proportional hazards models to evaluate their potential contributions to patient survival that are independent of *TP53* or *STAG2* mutation status (Additional file 2: Table S13). We performed the same analysis for 45 genes with potential different promoter usage in tumors with and without *TP53* mutations (Additional file 2: Table S14). Finally, we derived a multivariate Cox proportional hazards model including both *TP53* and *STAG2* mutation status, one candidate gene with differential promoter activity (*CALCB*), and five candidate gene isoforms (*CASZ1*, ENST00000496432; *RET*, ENST00000479913; *TEX40*, ENST00000328404; TNS1, ENST00000446903; and SLC27A6, ENST00000508645), and we followed this with backwards stepwise model selection. The final model (Figure 5b) revealed potential protective roles for one candidate transcript *TNS1* (ENST00000446903), a marginally protective role for candidate transcript *RET* (ENST00000479913), and a candidate transcript associated with a poor prognosis, *SLC27A6* (ENST00000508645).

*TNS1* encodes the well-studied protein Tensin 1, and is involved in several key aspects of cell function, including extracellular matrix formation, actin cytoskeleton formation, and signal transduction [64, 65]. More recently, the up-regulation of TNS1 in colorectal cancer was found to be associated with poor overall survival in patients [66], although previous studies have shown suppression of *TNS1* expression [67] is associated with metastatic cancers. Our results suggest that *TNS1* is a candidate prognostic indicator for EWS. Further studies are needed to draw more attention to the functional roles of these genes/transcripts in EWS progression.

## Discussion

Although epigenetic studies in disease models (cell lines, xenografts, and organoids) and in a limited number of primary tumor samples have demonstrated the oncogenic contributions of epigenetic deregulations to cancer initiation, progression, and response to treatment [48, 68, 69], genome-wide profiling of promoter activities by using standard approaches (e.g., ChIP-seq or CAGE) has not been carried out in large patient tumor cohorts, despite the continuous efforts of large epigenome consortia [45, 46, 70]. Our analyses of *MYCN*-amplified NBL tumors revealed both commonly active promoters and promoters that were active in some tumors but not in others. Moreover, these sample-specific active promoters are functionally important, as they drive the expression of several cancer consensus genes, including *MYC*. This observation is consistent with recent reports of heterogeneous enhancer activities of cell line–defined super-enhancers in primary gastric cancers [71], emphasizing the critical importance of deriving sample-specific epigenomic signatures. To bridge the gap between the extensive epigenomic resources in disease models and the limited ChIP-seq profiles of primary patient tumors, we developed MethylationToActivity (M2A), a deep-learning framework, to characterize the promoter activity landscape (both H3K4me3 and H3K27ac levels) in individual tumors by using DNAm data, which is the most extensively documented epigenetic modification for patient tumors and can be robustly and accurately profiled in FFPE archived retrospective samples. M2A demonstrated excellent performance across various tumor types, with accuracy comparable to that of ChIP-seq measurements of replicate samples from high-quality cohorts (Figure 3).

Although our framework was strictly trained on HM levels, the inferred promoter activity was highly correlated with the transcript-based gene expression levels quantified by RNA-seq (Additional file 2: Tables S5, S7 and S8). The correlation between gene expression and inferred promoter activities (mean *R*^2^ = 0.668 for ENCODE data, 0.722 for K562, 0.705 for GM12878, and 0.536 for H1-ESC) surpassed that with the state-of-art BPR model [42], which was developed for predicting gene expression levels from DNAm patterns (the best reported *R*^2^ values were 0.49 for K562, 0.37 for GM12878, and 0.25 for H1-ESC). Strikingly, the (indirect) predictive accuracy of M2A for gene expression across nine ENCODE cell lines (average *R*^2^ = 0.668) was comparable to the predictive accuracy of a model built on 11 HMs, one histone variant, and DNase I hypersensitivity [11]. Similarly, the accuracy of binary prediction of expressed genes (average AUC = 0.941 and 0.931 for the ENCODE and AML datasets, respectively) surpassed that of DeepChrome (average AUC = 0.80), a state-of-art deep-learning algorithm trained to predict expressed genes by using five core HMs (H3K4me3, H3K4me1, H3K36me3, H3K9me3, and H3K27me3) [12]. These results further validated our framework. Finally, both the M2A and BRP models suggest that it is insufficient to represent DNAm information by using a simple average methylation level in the promoter region. To properly reveal the regulatory roles of DNAm, we need to derive high-order features that capture spatial relations among CpG probes (or window-based derived features calculated from them) in promoters and in their vicinity. M2A uses the feature learning and selection characteristics of CNNs to achieve its exceptional performance, thus demonstrating the rich information content of DNAm signatures at both the genome-wide and local gene levels.

Analysis of the deep-learning framework revealed that M2A derives 1) high-level features from DNAm patterns that are common among different tumors and 2) tumor subtype–specific mapping functions from the mapping of high-level DNAm features to promoter activities in individual tumor subtypes by using transfer learning (when feasible). Although our deep-learning model cannot establish a causal relation between DNAm and promoter activities, these findings nevertheless shed light on both the general and tumor subtype–specific rules for interpreting DNAm patterns.

In evaluating our predictions, we found that several samples (the “poor performers”) showed abnormally low predictive accuracy with both the ENCODE and BLUEPRINT datasets. Investigations of the promoter H3K27ac levels revealed that the fraction of active promoters in these samples was substantially lower than in other samples. Furthermore, joint analyses with RNA-seq data from the matching samples indicated that 1) in contrast to the predictive accuracy of H3K27ac levels, the “poor performers” achieved comparable consistency between the M2A-inferred promoter activity and gene expression; 2) the “poor performers” showed significantly lower correlation between the measured promoter H3K27ac level and gene expression; 3) the correlation between the measured promoter H3K27ac level and gene expression was highly predictive of the accuracy of the H3K27ac prediction (Additional file 1: Figure S10),and 4) in ENCODE samples with replicates, the replicate with better consistency between H3K27ac and gene expression also showed significantly higher correlation between the actual and measured H3K27ac level (*P =* 0.0039, Wilcoxon signed rank test, Additional file 2: Table S15, an example of H3K27ac signal discrepancy between H1-ESC replicates shown in Additional file1: Figure S8d). Although we cannot unequivocally rule out the possibility that these “poor performers” share a distinct biological mechanism where promoter H3K27ac level is no longer a stronger predictor for gene activities, these results suggested that the “poor performers” could reflect the quality of the ChIP-seq results included in the test data. This observation emphasizes the value of conducting a preliminary analysis to evaluate the data quality before incorporating public data. It also suggests that M2A can provide a robust surrogate for promoter activities when the ChIP-seq experimental data is questionable.

Alternative promoter usage increases the transcriptomic diversity during normal tissue development and oncogenesis. Recent work demonstrated that an alternative promoter of *ERBB2* is predictive of a poor clinical outcome but that the canonical promoter shows no significant association with survival in patients with low-grade glioma [2]. Whereas earlier studies focused on identifying alternative promoter usage through CAGE or RNA-seq data, our research has shown that alternative promoter usage can be extensively studied by using DNAm profiles from diverse samples, including retrospective FFPE tumor samples, for which traditional approaches (CAGE, ChIP-seq, and RNA-seq) are technically challenging. Our analysis of a large EWS cohort (without matching RNA-seq data) revealed promoter activities for several specific isoforms that are independently associated with clinical outcomes, including a specific isoform of the *TNS1* gene (ENST00000446903). This demonstrates the importance of analyzing alternative promoter usage in epigenomic studies.

Finally, our analysis quantitatively emphasizes that promoter activity is one of the mechanisms that regulate the final transcriptional output: 1) both the observed and predicted promoter activities account for 50%–70% of the variation in gene expression, and 2) approximately 40% of the genes differentially expressed in ERMS and ARMS tumors show significant differences in their promoter activities. In addition to promoter activities, other epigenetic mechanisms, including enhancer activities, contribute substantially to gene regulation [72]. Recent work has demonstrated the roles of DNAm in regulating enhancer activities [73] and aberrant cancer-specific DNAm patterns in super-enhancers [74]. Consequently, we aim to expand our M2A framework to infer enhancer activities from DNAm patterns in the future.

## Conclusion

We have demonstrated that MethylationToActivity overcomes the unique challenges of systematically characterizing promoter activities from DNA methylation signatures. It achieved an accurate, robust and generalizable performance in various pediatric and adult cancers, including both solid and hematologic malignant neoplasms. MethylationToActivity will serve as a valuable tool to provide functional interpretation of DNAm deregulation, characterize promoter activity differences from DNAm patterns, and reveal alternate promoter usage in patient tumors, which will facilitate precision medicine by tailoring treatments based on both genetic variants and epigenetic deregulation.

## Methods

### Datasets

Five separate publicly available datasets were used in this study, including a pediatric NBL O-PDX dataset (N = 16) [40]; an RMS O-PDX dataset (N = 16) [42]; ENCODE datasets with matching H3K27ac and H3K4me3 histone mark ChIP-seq, RNA-seq, and WGBS experimental data (N = 9) [43]; a DCC BLUEPRINT AML dataset (N = 19) [44]; and a pediatric EWS dataset (N = 140) [19] (Additional file 2: Table S16). Of the 140 samples in the EWS cohort, only three had matching ChIP-seq and reduced-representation bisulfite sequencing (RRBS) data available; for the remaining 137 samples, only RRBS data was available. All other cohort datasets (i.e., the RMS, NBL, ENCODE, and AML datasets) contained matching H3K27ac and H3K4me3 profiles, along with RNA-seq and WGBS experimental data.

### Feature processing

All datasets were evaluated using GENCODE annotation definitions (www.gencodegenes.org/); the NBL, RMS, ENCODE, and EWS datasets were evaluated using GENECODE GRCh37.p13 (release 19), and the AML dataset was evaluated using GENCODE GRCh38.p13 (release 32). Promoter regions are defined as the TSS ± 1 kbp, where the TSS is defined as each unique transcript start position. To avoid using identical or near-identical promoter regions in training and baseline performance, only TSSs with promoter regions with less than 50% overlap were considered. Gene orientation was taken into account, and any promoters with overlying regions but opposite orientations were not considered as overlapping. Because of differences in sex amongst samples, all chromosome X and Y promoter regions were removed from consideration. This resulted in a total of 96,756 and 104,722 non-overlapping promoter regions from the annotation files GRCh37.p13 and GRCh38.p13, respectively. For gene expression analysis, we followed the definition in [11] and retained all 141,152 and 147,980 autosomal promoters for protein-coding genes from the annotation files GRCh37.p13 and GRCh38.p13, respectively.

M2A uses only one variable feature type: DNA methylation. For WGBS/RRBS data, the M-value was calculated as follows:

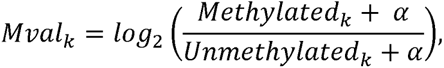

where *Methylated_k_* and *Unmethylated_k_* correspond to the number of methylated and unmethylated reads of the *k^th^* CpG site, respectively. By default, the offset was set to 0.5, and a global M-value threshold was set to a maximum value of log_2_65 and a minimum value of 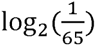. CpG sites with coverage of less than five reads were removed.

M2A uses a promoter region–based windowed approach, comprising 20 windows of two sizes (250 bp and 2.5 kbp) and a step size equal to the window size, centered on a given TSS. For instance, ***W_ij_*** = {*W_i_*_1_,*W_i_*2,*W_i_*3 … *W_i_*_20_} is the vector of windows where *i* represents a particular TSS and ***j*** represents a particular window corresponding with the ***i^th^*** TSS. This means that *W_i_*_10_ and *W_i_*_11_ represent the windows immediately downstream and upstream of the ***i^th^*** TSS. Therefore,

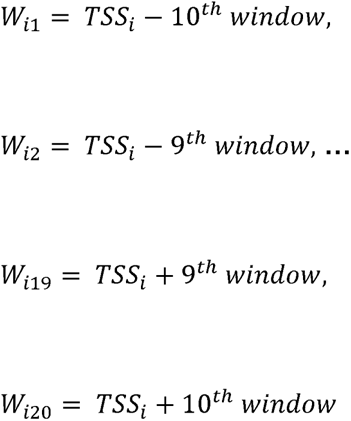

Each feature was calculated in this manner. The DNA methylation features, including the windowed M-value mean, variance, and the fraction of the SSD of M-values (FSSD), were calculated and represented by the feature vectors ***Mave_i_***, ***Mvar***_i_ and ***Mfssd***_i_. Therefore, the features for a particular window, denoted as *W_ij_*, would be calculated as follows:

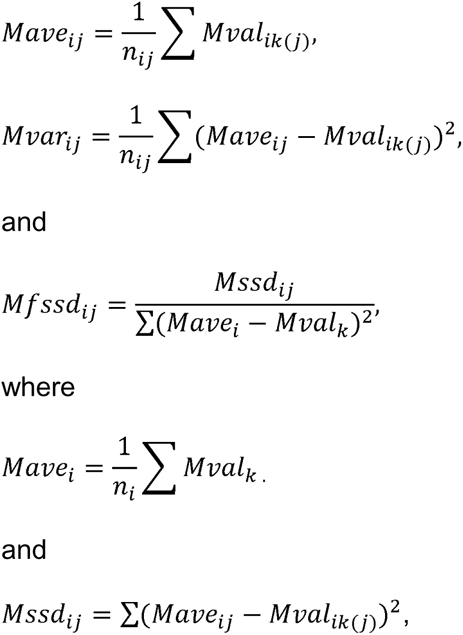

Here, *i* represents the promoter, *j* represents a specific window for a particular promoter, and *Mval_k_* represents the Mval for individual CpGs in a region where ***Mval_k(j)_*** is the Mval for an individual CpG in a specific window. Each feature was interleaved by window size to provide model input wherein each window contained a number of “channels” equal to the number of features, with the feature array shape being (*N*, 2, 20, 3), where *N* represents the total number of TSSs in a sample, 2 represents the number of window sizes (250 bp and 2.5 kbp), 20 represents the number of windows, and 4 represents the number of features per window. All features were scaled from 0.1 to 1 (using MinMaxScaler with default values from sklearn version 0.22); in instances where windows overlapped regions without methylation data, resulting in NaNs (such as chromosomal boundaries, telomeric regions, and centromeric regions), these feature values were marked as 0.

### Calculating histone modification enrichment

The response variable was calculated for each non-overlapping promoter region, a 2000-bp region centered on each TSS. Histone modification enrichment (*HM*) for the *i^th^* promoter region is calculated as follows:

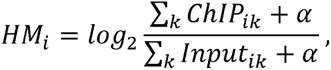

where ∑*_k_ChlP_ik_* represents the sum of either the H3K27ac or the H3K4me3 read signal mapped to the promoter region at each position, ∑*_k_input_ik_* represents the sum of the control read signal mapped to the promoter region, and α represents the 25^th^ percentile of the ∑*_k_input_ik_* calculated for a given sample.

### M2A topology

M2A is a machine learning framework that leverages canonical deep-learning strategies, including convolutional neural network (CNN) and fully connected (FC) layers. Each layer employs a LeakyReLU (alpha = 0.1) and a kernel constraint by L2-normalization using maxnorm(3). CNN layers are two-dimensional, with zero padding, a stepsize of (1,1), and a kernel size of (1, 3) to maintain feature space and prevent convolutions across features from different window sizes. To test the efficacy of this approach, we compared the performance of a traditional artificial neural network (ANN) consisting of two FC layers versus three CNN layers in addition to two FC layers. During transfer learning, weights corresponding to each of the three CNN layers of the six–NBL O-PDX M2A model trained previously were frozen; only the weights corresponding to the two FC layers were optimized. A summary of all model topologies and parameters can be found in Additional file 2: Table S17. To train and test each model topology, we used Keras (v2.2.4) and Tensorflow (v2.1.0) in Python 3.6.5.

### M2A parameter tuning

Parameters such as the window size, batch size, and kernel size were optimized using the validation NBL data set (n=10). Each parameter configuration was tested holding all other parameters constant, and models with a numerical performance advantage were chosen. For batch size, three configurations were tested (64, 128, 256). Four kernel size configurations ([1,2], [1,3], [1,4], [1,5]) were tested. and window configurations ([100bp, 1000bp], [250bp, 2500bp], [500bp, 5000bp]) were considered. Due to >50% uninformative features in the 100bp window resolution, only the (250bp, 2500bp) model and (500bp, 5000bp) model performances were compared (Additional file 1: Figure S13; Additional file 2: Table S18).

### Training M2A

The core M2A model (without transfer) training set consisted of six O-PDX samples from the 16-sample NBL cohort (Additional file 2: Table S19); we trained separate models for H3K27ac and H3K4me3 HMs with the same WGBS features as the input. After the base models were trained, transfer learning was employed for three separate datasets, namely the RMS, AML, and EWS datasets. Each transfer learning model was trained using one sample from the cohort, for a total of N models, where N equals the number of samples in the cohort. For the RMS and EWS cohorts, an ensemble approach was used, whereby an averaged prediction from N−1 models was generated after transfer learning with each sample. The same approach was used with the AML cohort, except that only samples with *R*^2^ ≥ 0.60 between FPKM and H3K27ac were used for transfer learning.

For each training scheme, the same parameters were used, including an 80/20 training/validation split and a batch size of 64 (Additional file 2: Table S17). All sample input was randomized before training. The Keras implementation of adadelta (default parameters) minimizing the mean squared error (MSE) was used to optimize M2A. To prevent overtraining, the EarlyStopping method was employed by monitoring validation loss for 10 epochs without at least a minimal gain in performance (min_delta = 0.0001) for a maximum of 80 epochs. In no case was the maximum number of epochs reached.

### Determining promoter diversity in NBL models

Promoter region–based H3K27ac distributions clearly show a bimodal distribution (see Additional file 1: Figure S1); therefore, to determine class occupancy (active versus inactive), a Gaussian mixture model (GaussianMixture from sklearn version 0.22 with n_components = 2) was applied for each individual sample. To determine the percentage of differentially active promoters among all active promoters, we used a pairwise comparison approach for all samples. Cancer consensus genes were downloaded from COSMIC (https://cancer.sanger.ac.uk/census) (accessed on February 1, 2020). To avoid artificially inflated values from genes with multiple TSSs, only cancer consensus genes with a single TSS according to GENCODE GRCh37.p13 (release 19) definitions were considered.

### Evaluating M2A performance

When determining prediction performance, two primary metrics were considered, namely the *R*^2^ and the root mean squared error (RMSE). To measure the accuracy of M2A in predicting expressed genes, we calculated the AUC-ROC by using roc_curve and the precision-recall curve AUC by using average_precision_score from sklearn 0.22. Paired analyses were tested for statistical significance by using a Wilcoxon signed-rank test (R v3.4.1). To determine outliers, a median-based method was implemented using the “outlier” function in the R package GmAMisc [55]. To ensure that low-mappability regions were not a confounding factor, we used the wgEncodeCrgMApabilityAlign100mer.bw file downloaded from the UCSC Genome Browser (http://genome.ucsc.edu/cgi-bin/hgFileUi?db=hg19&g=wgEncodeMapability). For GRCh38.p13 annotations, we used liftOver from UCSC (http://hgdownload.cse.ucsc.edu/admin/exe/linux.x86_64/) to convert the mappability track to GRCh38. For all performance-related analyses, the average value of this track within each non-overlapping 2000-bp promoter region was calculated, and promoter regions with mappability > 0.75 were retained for evaluation.

In analyses between H3K27ac levels (actual or predicted) and gene expression, we followed the gene filtering steps described in [11]: all autosomal protein-coding gene promoters were considered and for genes with multiple promoters, we use the maximum promoter activity to represent the gene.

### Baseline models

Multivariate adaptive regression splines (MARS) was implemented by “earth” package in R (all default values were used), and the random forest baseline model was implemented by the “sklearn.ensemble.RandomForestRegressor” package in python 3.7.0 (max_features=’sqrt’). For comparison purposes, each model tested used identical feature input to M2A.

### M2A captures the impact of DMRs on promoter region activity

The lists of genes that are differentially expressed in ERMS and ARMS samples and genes with DMRs were previously reported in reference [42].

### Alternative promoter usage analyses

To infer alternative promoter usage that was specific to the RMS subtypes ARMS and ERMS, we first delineated the “active” vs. “inactive” promoters in a subtype (ERMS or ARMS) by applying a threshold Of mean(H3K27ac) > 1 for the average samples in the subtype. Next, primary promoters from multi-promoter genes were determined by the average H3K27ac level within a specific subtype, and the promoter with the maximum activity from a given gene was counted as the primary promoter. The differences in promoter usage between two subtypes was defined as the difference between the activity sum of the primary promoters and the activity sum of the secondary promoters in the two subtypes.

### Analysis of alternative promoter usage in EWS

In the same manner as the RMS subtype analysis, EWS alternative promoter usage was determined in two patient sample groups: *TP53* mutant and *TP53* wild-type groups (for sample status, see Additional file 2: Table S18). Candidate genes were identified as active genes (H3K27ac ≥ 1 in at least one group) with potential differential promoter activities (absolute difference ≥ 1) in the *TP53* wild-type and *TP53* mutant groups. Candidate genes with alternative promoter usage were identified as genes that used different active primary promoters in the wild-type and mutant groups and had an average promoter usage difference of at least 0.4 between groups. Both alternative promoters and differentially active promoters were considered in prognostic analyses (univariate screening incorporating both *TP53* and *STAG2* mutation status, followed by a multivariate analysis) using Cox proportional hazard models (R 3.4.1). The final model was derived from backward stepwise selection from a Cox proportional hazards model including *TP53* and *STAG2* mutation status and all potential markers (all genes or promoters with an FDR < 0.05 in the univariate analysis).

### Analysis of M2A feature input and extracted features

To determine the merit of a CNN-based approach, each feature average for a particular window position and window size were plotted with a 95% confidence interval. The plotted feature distribution was calculated from all M2A vanilla model training data input (NBL, N=6), and stratified by promoter status. Promoter status was determined by class occupancy of both H3K27ac and H3K4me3, (active versus inactive), where (H3K27ac=active), (H3K27ac=inactive, H3K4me3=active), and (H3K27ac=inactive, H3K4me3=inactive), represents active, poised, and inactive promoters, respectively. Class occupancy was determined by applying a Gaussian mixture model (GaussianMixture from sklearn version 0.22 with n_components = 2).

The efficacy of the CNN-based feature extraction was tested by 1) training sample input feature predictive performance as compared to CNN-extracted feature performance, calculated by Pearson’s *R^2^* for all features across the entire training set (each window size considered separately; NBL, N=6) and 2) The performance (Pearson’s *R^2^*) of the best performing feature identified in 1) when applied to each sample in the validation set (NBL, N=10; Additional File 1: Figure S3).

### M2A code availability

The latest M2A models, feature generation, prediction pipeline, and a Docker image of the M2A environment pre-loaded are available for download at https://github.com/chenlab-sj/M2A. Additionally, source code with detailed instructions for transfer learning using the M2A model with input samples from other domains is available. The cloud-based implementation of M2A is available to anyone with a (free) St. Jude Cloud account (https://platform.stjude.cloud/workflows/methylation-to-activity).

## Declarations

### Ethics approval and consent to participate

Not applicable

### Consent for publication

Not applicable

### Availability of data and materials

The datasets generated and/or analyzed during the current study are publicly available, as detailed in Additional File 2: Table S16.

### Authors’ contributions

**JW**: Methodology, software, data analysis, result interpretation, writing - original draft, writing - review & editing. **BX**: Resources, data curation. **DP**: Software, resources. **AT**: Software, resources. **CL**: Result interpretation, writing – review & editing. **JY**: Result interpretation, writing – review & editing. **XC**: Conceptualization, methodology, data analysis, result interpretation, writing - review & editing, supervision.

## Acknowledgements

We thank Keith A. Laycock, PhD, ELS, for editing the manuscript.

## Additional Files

### Additional file 1 (file type .PDF, 7 MB)

**Figure S1. NBL sample H3K27ac promoter distribution.** A comparison of promoter H3K27ac enrichment in *MYCN* amplified (MNA) NBL cell line and O-PDX samples shows a clear bimodal distribution, delineating “active” and “non-active” promoters.

**Figure S2. DNAm input feature pattern analysis.** The DNAm features show promoter-status-specific patterns between active, poised, and inactive promoters, emphasizing the utility of positional-based relationships in a windowed DNAm feature approach. For both 250 bp and 2,500 bp window sizes, the average scaled input feature was plotted in relationship to the TSS (ribbons represent the 95% confidence interval), stratified by the promoter status. Promoter status was determined by class occupancy of both H3K27ac and H3K4me3, either “active” or “inactive”, where (H3K27ac=active), (H3K27ac=inactive, H3K4me3=active), and (H3K27ac=inactive, H3K4me3=inactive), represents active, poised, and inactive promoters, respectively.

**Figure S3. Feature performance comparison: Input features vs CNN mapped features.** Using all samples included in training the vanilla M2A model (NBL, N=6), the individual feature performance (as determined by Pearson’s *R^2^* between the feature and the response variable, H3K27ac) for each feature was plotted, comparing the distribution of performances between raw input training feature and the CNN mapped features at a particular window size (250 bp or 2,500 bp). The best feature from this analysis for each window size and feature type (input or CNN mapped) was used to determine Pearson’s *R^2^* with H3K27ac from each sample in the NBL validation set (N=10).

**Figure S4. M2A prediction generalizability analysis. (a)** A comparison between performance of the M2A model (*R^2^* of observed H3K27ac promoter levels and predicted levels in the test sample) with the surrogate model (represented by the highest *R^2^* of observed H3K27ac promoter levels in the test sample and the observed H3K27ac promoter levels in any training sample). M2A extracts generalizable features capable of out-performing the surrogate model in both the NBL validation set and the RMS test set, further highlighted in **(b),** a comparison of surrogate model and M2A model performance using promoters from DE genes between NBL and RMS.

**Figure S5. M2A with transfer learning outperforms a vanilla M2A model of the same cancer type.** An M2A model with transfer learning (initially trained with six NBL O-PDX samples and then transferred with a single RMS sample) consistently outperforms an M2A model trained with a single RMS sample.

**Figure S6. Signal-to-noise analysis of ENCODE and NBL datasets.** Comparison at the observed H3K27ac promoter level **(a)** and the H3K4me3 **(b)** promoter levels revealed different signal-to-noise profiles between the ENCODE dataset and the NBL datasets, which results in a highly correlated prediction with larger RMSE values.

**Figure S7. M2A ENCODE cohort performance.** The distribution of M2A prediction performance (*R^2^*), shows that M2A accurately infers both H3K27ac and H4K4me3 promoter levels in the publicly available ENCODE dataset.

**Figure S8. Analysis of outlier H1-ESC. (a)** When inferring H3K27ac promoter levels, M2A was substantially less accurate in the ENCODE sample H1-ESC, which is an outlier in the ENCODE cohort. **(b, c)** M2A-predicted H3K27ac promoter levels **(b)** are more consistent with, and more predictive of, H1-ESC gene expression than are the actual observed H1-ESC H3K27ac promoter levels **(c)**. (**d**) The hypomethylated region surrounding the promoters of genes *PSMA7* and *SS18L1* (often indicative of H3K27ac enrichment) showed inconsistent H3K27ac levels between H1-ESC ChIP-seq replicates from ENCODE.

**Figure S9. Predicting gene expression in the ENCODE dataset. (a–i)** The indirect ability of M2A to predict gene expression (i.e., expressed vs. not expressed) on the basis of both M2A-predicted and observed H3K27ac promoter levels for each sample was determined by comparing the AUCs of the receiver operating characteristic (ROC) curves.

**Figure S10. Consistency of gene expression and H3K27ac promoter levels in the AML cohort.** The consistency, as determined by Pearson’s *R*^2^, between the observed values for gene expression and the H3K27ac promoter levels is remarkably predictive of the performance M2A in predicting H3K27ac promoter levels in samples from the AML cohort.

**Figure S11. M2A accurately determines subtype differences between embryonal and alveolar RMS. (a)** The promoter activities of single-promoter, differentially expressed genes in the RMS subtypes ERMS and ARMS are accurately inferred by an M2A base model (trained with six O-PDX NBL samples). **(b)** The predictive performance of M2A is further boosted by transfer learning with one RMS sample. **(c, d)** The performance of M2A declines slightly when the model is applied to all promoters of differentially expressed genes **(c)**, but it recovers when an M2A model with transfer learning with only one RMS training sample is applied **(d)**.

**Figure S12. Kaplan–Meier log-rank analysis by mutation status in EWS.** The prognostic ability of **(a)** *TP53* or **(b)** *STAG2* mutation status in the EWS cohort was determined by the log-rank test and visualized using the Kaplan–Meier survival curve. The *STAG2* mutation status showed no significant difference in overall survivability, thus only *TP53* mutation status was considered when forming the univariate and multivariate Cox proportional hazards model.

**Figure S13. CpG distribution by window relative to the TSS.** To achieve feature input that is informative to the model, M2A window size selection was partially based on the number of CpGs captured by window size. Each analysis consists of 20 windows surrounding each TSS at a particular window size, representing the theoretical CpG input to M2A for that particular resolution. Three different window configurations were considered, comprised of two window sizes: 1) 100 bp and 1,000 bp, 2) 250 bp and 2,500 bp, and 3) 500 bp and 5,000 bp. Due to NaNs in feature windows calculated with fewer than 2 CpGs, the [100 bp, 1000 bp] model was removed from consideration (> 50% NaNs).

### Additional file 2 (file type: .XLSX, 179 KB)

Table S1: H3K27ac active cancer consensus genes in 3 NBL cell lines, and 3 NBL O–PDX samples.

Table S2: Baseline models vs. vanilla M2A predictive performance comparison.

Table S3: M2A predictive performance in NBL cell line samples.

Table S4: Observed H3K27ac and H3K4me3 ENCODE replicate consistencies.

Table S5: M2A predictive performance in RMS O–PDX samples.

Table S6: M2A RMS transfer model predictive performance in RMS O–PDX samples.

Table S7: M2A predictive performance in ENCODE dataset.

Table S8: M2A predictive performance in AML samples.

Table S9: ERMS vs. ARMS DMRs and associated genes (overexpressed in ERMS).

Table S10: ERMS vs. ARMS DMRs and associated genes (overexpressed in ARMS).

Table S11: M2A alternate promoter usage predictive performance between ARMS and ERMS samples.

Table S12: M2A predictive performance in EWS samples, before and after transfer.

Table S13: A univariate survival analysis of differential H3K27ac promoter activity between TP53 mutant and TP53 wild–type EWS tumors.

Table S14: A univariate survival analysis of alternate promoter usage between TP53 mutant and TP53 wild–type EWS tumors.

Table S15: ENCODE H3K27ac replicate consistency with gene expression.

Table S16: Dataset availability.

Table S17: M2A model topologies.

Table S18: Parameter tuning: mean performance in the NBL validation set (*R^2^*)

Table S19: Sample summary information.

## References

1. Davuluri RV, Suzuki Y, Sugano S, Plass C, Huang TH: The functional consequences of alternative promoter use in mammalian genomes. Trends Genet 2008, 24:167–177.

2. Demircioglu D, Cukuroglu E, Kindermans M, Nandi T, Calabrese C, Fonseca NA, Kahles A, Lehmann KV, Stegle O, Brazma A, et al: A Pan-cancer Transcriptome Analysis Reveals Pervasive Regulation through Alternative Promoters. Cell 2019, 178:1465–1477 e1417.

3. Qamra A, Xing M, Padmanabhan N, Kwok JJT, Zhang S, Xu C, Leong YS, Lee Lim AP, Tang Q, Ooi WF, et al: Epigenomic Promoter Alterations Amplify Gene Isoform and Immunogenic Diversity in Gastric Adenocarcinoma. Cancer Discov 2017, 7:630–651.

4. Sotillo E, Barrett DM, Black KL, Bagashev A, Oldridge D, Wu G, Sussman R, Lanauze C, Ruella M, Gazzara MR, et al: Convergence of Acquired Mutations and Alternative Splicing of CD19 Enables Resistance to CART-19 Immunotherapy. Cancer Discov 2015, 5:1282–1295.

5. Ma X, Liu Y, Liu Y, Alexandrov LB, Edmonson MN, Gawad C, Zhou X, Li Y, Rusch MC, Easton J, et al: Pan-cancer genome and transcriptome analyses of 1,699 paediatric leukaemias and solid tumours. Nature 2018, 555:371–376.

6. Grobner SN, Worst BC, Weischenfeldt J, Buchhalter I, Kleinheinz K, Rudneva VA, Johann PD, Balasubramanian GP, Segura-Wang M, Brabetz S, et al: The landscape of genomic alterations across childhood cancers. Nature 2018, 555:321–327.

7. Huether R, Dong L, Chen X, Wu G, Parker M, Wei L, Ma J, Edmonson MN, Hedlund EK, Rusch MC, et al: The landscape of somatic mutations in epigenetic regulators across 1,000 paediatric cancer genomes. Nat Commun 2014, 5:3630.

8. Andersson R, Sandelin A: Determinants of enhancer and promoter activities of regulatory elements. Nat Rev Genet 2020, 21:71–87.

9. Kelley DZ, Flam EL, Izumchenko E, Danilova LV, Wulf HA, Guo T, Singman DA, Afsari B, Skaist AM, Considine M, et al: Integrated Analysis of Whole-Genome ChIP-Seq and RNA-Seq Data of Primary Head and Neck Tumor Samples Associates HPV Integration Sites with Open Chromatin Marks. Cancer Res 2017, 77:6538–6550.

10. Karlic R, Chung HR, Lasserre J, Vlahovicek K, Vingron M: Histone modification levels are predictive for gene expression. Proc Natl Acad Sci U S A 2010, 107:2926–2931.

11. Dong X, Greven MC, Kundaje A, Djebali S, Brown JB, Cheng C, Gingeras TR, Gerstein M, Guigo R, Birney E, Weng Z: Modeling gene expression using chromatin features in various cellular contexts. Genome Biol 2012, 13:R53.

12. Singh R, Lanchantin J, Robins G, Qi Y: DeepChrome: deep-learning for predicting gene expression from histone modifications. Bioinformatics 2016, 32:i639–i648.

13. Kagohara LT, Stein-O’Brien GL, Kelley D, Flam E, Wick HC, Danilova LV, Easwaran H, Favorov AV, Qian J, Gaykalova DA, Fertig EJ: Epigenetic regulation of gene expression in cancer: techniques, resources and analysis. Brief Funct Genomics 2018, 17:49–63.

14. Zhang P, Lehmann BD, Shyr Y, Guo Y: The Utilization of Formalin Fixed-Paraffin-Embedded Specimens in High Throughput Genomic Studies. Int J Genomics 2017, 2017:1926304.

15. Moran S, Vizoso M, Martinez-Cardus A, Gomez A, Matias-Guiu X, Chiavenna SM, Fernandez AG, Esteller M: Validation of DNA methylation profiling in formalin-fixed paraffin-embedded samples using the Infinium HumanMethylation450 Microarray. Epigenetics 2014, 9:829–833.

16. de Ruijter TC, de Hoon JP, Slaats J, de Vries B, Janssen MJ, van Wezel T, Aarts MJ, van Engeland M, Tjan-Heijnen VC, Van Neste L, Veeck J: Formalin-fixed, paraffin-embedded (FFPE) tissue epigenomics using Infinium HumanMethylation450 BeadChip assays. Lab Invest 2015, 95:833–842.

17. Gu H, Bock C, Mikkelsen TS, Jager N, Smith ZD, Tomazou E, Gnirke A, Lander ES, Meissner A: Genome-scale DNA methylation mapping of clinical samples at single-nucleotide resolution. Nat Methods 2010, 7:133–136.

18. Ziller MJ, Gu H, Muller F, Donaghey J, Tsai LT, Kohlbacher O, De Jager PL, Rosen ED, Bennett DA, Bernstein BE, et al: Charting a dynamic DNA methylation landscape of the human genome. Nature 2013, 500:477–481.

19. Charlet J, Duymich CE, Lay FD, Mundbjerg K, Dalsgaard Sorensen K, Liang G, Jones PA: Bivalent Regions of Cytosine Methylation and H3K27 Acetylation Suggest an Active Role for DNA Methylation at Enhancers. Mol Cell 2016, 62:422–431.

20. Onuchic V, Lurie E, Carrero I, Pawliczek P, Patel RY, Rozowsky J, Galeev T, Huang Z, Altshuler RC, Zhang Z, et al: Allele-specific epigenome maps reveal sequence-dependent stochastic switching at regulatory loci. Science 2018, 361.

21. Stadler MB, Murr R, Burger L, Ivanek R, Lienert F, Scholer A, van Nimwegen E, Wirbelauer C, Oakeley EJ, Gaidatzis D, et al: DNA-binding factors shape the mouse methylome at distal regulatory regions. Nature 2011, 480:490–495.

22. Sheffield NC, Pierron G, Klughammer J, Datlinger P, Schonegger A, Schuster M, Hadler J, Surdez D, Guillemot D, Lapouble E, et al: DNA methylation heterogeneity defines a disease spectrum in Ewing sarcoma. Nat Med 2017, 23:386–395.

23. Zhu H, Wang G, Qian J: Transcription factors as readers and effectors of DNA methylation. Nature Reviews Genetics 2016, 17:551–565.

24. Kondo Y: Epigenetic cross-talk between DNA methylation and histone modifications in human cancers. Yonsei Med J 2009, 50:455–463.

25. Rothbart SB, Strahl BD: Interpreting the language of histone and DNA modifications. Biochim Biophys Acta 2014, 1839:627–643.

26. Hashimshony T, Zhang J, Keshet I, Bustin M, Cedar H: The role of DNA methylation in setting up chromatin structure during development. Nat Genet 2003, 34:187–192.

27. Moore LD, Le T, Fan G: DNA methylation and its basic function. Neuropsychopharmacology 2013, 38:23–38.

28. Jenkinson G, Pujadas E, Goutsias J, Feinberg AP: Potential energy landscapes identify the information-theoretic nature of the epigenome. Nat Genet 2017, 49:719–729.

29. Fortin JP, Hansen KD: Reconstructing A/B compartments as revealed by Hi-C using long-range correlations in epigenetic data. Genome Biol 2015, 16:180.

30. Simmonds P, Loomis E, Curry E: DNA methylation-based chromatin compartments and ChIP-seq profiles reveal transcriptional drivers of prostate carcinogenesis. Genome Med 2017, 9:54.

31. Mack SC, Witt H, Piro RM, Gu L, Zuyderduyn S, Stutz AM, Wang X, Gallo M, Garzia L, Zayne K, et al: Epigenomic alterations define lethal CIMP-positive ependymomas of infancy. Nature 2014, 506:445–450.

32. Teodoridis JM, Hardie C, Brown R: CpG island methylator phenotype (CIMP) in cancer: causes and implications. Cancer Lett 2008, 268:177–186.

33. Capper D, Jones DTW, Sill M, Hovestadt V, Schrimpf D, Sturm D, Koelsche C, Sahm F, Chavez L, Reuss DE, et al: DNA methylation-based classification of central nervous system tumours. Nature 2018, 555:469–474.

34. Perez E, Capper D: Invited Review: DNA methylation-based classification of paediatric brain tumours. Neuropathol Appl Neurobiol 2020.

35. Strahl BD, Allis CD: The language of covalent histone modifications. Nature 2000, 403:41–45.

36. Musselman CA, Lalonde ME, Cote J, Kutateladze TG: Perceiving the epigenetic landscape through histone readers. Nat Struct Mol Biol 2012, 19:1218–1227.

37. Spainhour JC, Lim HS, Yi SV, Qiu P: Correlation Patterns Between DNA Methylation and Gene Expression in The Cancer Genome Atlas. Cancer Inform 2019, 18:1176935119828776.

38. Baylin SB: DNA methylation and gene silencing in cancer. Nature Clinical Practice Oncology 2005, 2:S4–S11.

39. Jones PA: Functions of DNA methylation: islands, start sites, gene bodies and beyond. Nature Reviews Genetics 2012, 13:484–492.

40. Lay FD, Liu Y, Kelly TK, Witt H, Farnham PJ, Jones PA, Berman BP: The role of DNA methylation in directing the functional organization of the cancer epigenome. Genome Research 2015, 25:467–477.

41. Kagohara LT, Stein-O’Brien GL, Kelley D, Flam E, Wick HC, Danilova LV, Easwaran H, Favorov AV, Qian J, Gaykalova DA, Fertig EJ: Epigenetic regulation of gene expression in cancer: techniques, resources and analysis. Briefings in Functional Genomics 2018, 17:49–63.

42. Kapourani CA, Sanguinetti G: Higher order methylation features for clustering and prediction in epigenomic studies. Bioinformatics 2016, 32:i405–i412.

43. Zeineldin M, Federico S, Chen X, Fan Y, Xu B, Stewart E, Zhou X, Jeon J, Griffiths L, Nguyen R, et al: MYCN amplification and ATRX mutations are incompatible in neuroblastoma. Nat Commun 2020, 11:913.

44. Stewart E, McEvoy J, Wang H, Chen X, Honnell V, Ocarz M, Gordon B, Dapper J, Blankenship K, Yang Y, et al: Identification of Therapeutic Targets in Rhabdomyosarcoma through Integrated Genomic, Epigenomic, and Proteomic Analyses. Cancer Cell 2018.

45. Consortium EP: An integrated encyclopedia of DNA elements in the human genome. Nature 2012, 489:57–74.

46. Stunnenberg HG, International Human Epigenome C, Hirst M: The International Human Epigenome Consortium: A Blueprint for Scientific Collaboration and Discovery. Cell 2016, 167:1145–1149.

47. Singh AA, Schuurman K, Nevedomskaya E, Stelloo S, Linder S, Droog M, Kim Y, Sanders J, van der Poel H, Bergman AM, et al: Optimized ChIP-seq method facilitates transcription factor profiling in human tumors. Life Sci Alliance 2019, 2:e201800115.

48. Seligson DB, Horvath S, Shi T, Yu H, Tze S, Grunstein M, Kurdistani SK: Global histone modification patterns predict risk of prostate cancer recurrence. Nature 2005, 435:1262–1266.

49. Chen X, Stewart E, Shelat AA, Qu C, Bahrami A, Hatley M, Wu G, Bradley C, McEvoy J, Pappo A, et al: Targeting oxidative stress in embryonal rhabdomyosarcoma. Cancer Cell 2013, 24:710–724.

50. Stewart E, Shelat A, Bradley C, Chen X, Federico S, Thiagarajan S, Shirinifard A, Bahrami A, Pappo A, Qu C, et al: Development and characterization of a human orthotopic neuroblastoma xenograft. Dev Biol 2015, 407:344–355.

51. Stewart E, Federico SM, Chen X, Shelat AA, Bradley C, Gordon B, Karlstrom A, Twarog NR, Clay MR, Bahrami A, et al: Orthotopic patient-derived xenografts of paediatric solid tumours. Nature 2017, 549:96–100.

52. Murphy AJ, Chen X, Pinto EM, Williams JS, Clay MR, Pounds SB, Cao X, Shi L, Lin T, Neale G, et al: Forty-five patient-derived xenografts capture the clinical and biological heterogeneity of Wilms tumor. Nat Commun 2019, 10:5806.

53. Angermueller C, Lee HJ, Reik W, Stegle O: DeepCpG: accurate prediction of single-cell DNA methylation states using deep learning. Genome Biol 2017, 18:67.

54. Barredo Arrieta A, Díaz-Rodríguez N, Del Ser J, Bennetot A, Tabik S, Barbado A, Garcia S, Gil-Lopez S, Molina D, Benjamins R, et al: Explainable Artificial Intelligence (XAI): Concepts, taxonomies, opportunities and challenges toward responsible AI. Information Fusion 2020, 58:82–115.

55. Wilcox RR: The Normal Curve and Outlier Detection. In Fundamentals of Modern Statistical Methods. 2001: 31–47

56. Golob JL, Kumar RM, Guenther MG, Pabon LM, Pratt GA, Loring JF, Laurent LC, Young RA, Murry CE: Evidence that gene activation and silencing during stem cell differentiation requires a transcriptionally paused intermediate state. PLoS One 2011, 6:e22416.

57. Berdasco M, Esteller M: Clinical epigenetics: seizing opportunities for translation. Nat Rev Genet 2019, 20:109–127.

58. Hinoue T, Weisenberger DJ, Lange CPE, Shen H, Byun HM, Van Den Berg D, Malik S, Pan F, Noushmehr H, van Dijk CM, et al: Genome-scale analysis of aberrant DNA methylation in colorectal cancer. Genome Research 2011, 22:271–282.

59. Cao L, Yu Y, Bilke S, Walker RL, Mayeenuddin LH, Azorsa DO, Yang F, Pineda M, Helman LJ, Meltzer PS: Genome-wide identification of PAX3-FKHR binding sites in rhabdomyosarcoma reveals candidate target genes important for development and cancer. Cancer research 2010, 70:6497–6508.

60. Laé M, Ahn E, Mercado G, Chuai S, Edgar M, Pawel B, Olshen A, Barr F, Ladanyi M: Global gene expression profiling of PAX-FKHR fusion-positive alveolar and PAX-FKHR fusion-negative embryonal rhabdomyosarcomas. The Journal of Pathology 2007, 212:143–151.

61. Marshall AD, Grosveld GC: Alveolar rhabdomyosarcoma – The molecular drivers of PAX3/7-FOXO1-induced tumorigenesis. Skeletal Muscle 2012, 2:25.

62. Honda T, Inui M: PDZRN3 regulates differentiation of myoblasts into myotubes through transcriptional and posttranslational control of Id2. Journal of Cellular Physiology 2019, 234:2963–2972.

63. Tirode F, Surdez D, Ma X, Parker M, Le Deley MC, Bahrami A, Zhang Z, Lapouble E, Grossetete-Lalami S, Rusch M, et al: Genomic Landscape of Ewing Sarcoma Defines an Aggressive Subtype with Co-Association of STAG2 and TP53 Mutations. Cancer Discov 2014, 4:1342–1353.

64. Lo SH, An Q, Bao S, Wong WK, Liu Y, Janmey PA, Hartwig JH, Chen LB: Molecular cloning of chick cardiac muscle tensin. Full-length cDNA sequence, expression, and characterization. J Biol Chem 1994, 269:22310–22319.

65. Chen H, Duncan IC, Bozorgchami H, Lo SH: Tensin1 and a previously undocumented family member, tensin2, positively regulate cell migration. Proc Natl Acad Sci U S A 2002, 99:733–738.

66. Zhou H, Zhang Y, Wu L, Xie W, Li L, Yuan Y, Chen Y, Lin Y, He X: Elevated transgelin/TNS1 expression is a potential biomarker in human colorectal cancer. Oncotarget 2018, 9:1107–1113.

67. Hall EH, Daugherty AE, Choi CK, Horwitz AF, Brautigan DL: Tensin1 requires protein phosphatase-1alpha in addition to RhoGAP DLC-1 to control cell polarization, migration, and invasion. J Biol Chem 2009, 284:34713–34722.

68. Shen H, Laird PW: Interplay between the cancer genome and epigenome. Cell 2013, 153:38–55.

69. De Craene B, Berx G: Regulatory networks defining EMT during cancer initiation and progression. Nat Rev Cancer 2013, 13:97–110.

70. Roadmap Epigenomics C, Kundaje A, Meuleman W, Ernst J, Bilenky M, Yen A, Heravi-Moussavi A, Kheradpour P, Zhang Z, Wang J, et al: Integrative analysis of 111 reference human epigenomes. Nature 2015, 518:317–330.

71. Ooi WF, Xing M, Xu C, Yao X, Ramlee MK, Lim MC, Cao F, Lim K, Babu D, Poon LF, et al: Epigenomic profiling of primary gastric adenocarcinoma reveals super-enhancer heterogeneity. Nat Commun 2016, 7:12983.

72. Pennacchio LA, Bickmore W, Dean A, Nobrega MA, Bejerano G: Enhancers: five essential questions. Nat Rev Genet 2013, 14:288–295.

73. Weigel C, Chaisaingmongkol J, Assenov Y, Kuhmann C, Winkler V, Santi I, Bogatyrova O, Kaucher S, Bermejo JL, Leung SY, et al: DNA methylation at an enhancer of the three prime repair exonuclease 2 gene (TREX2) is linked to gene expression and survival in laryngeal cancer. Clin Epigenetics 2019, 11:67.

74. Heyn H, Vidal E, Ferreira HJ, Vizoso M, Sayols S, Gomez A, Moran S, Boque-Sastre R, Guil S, Martinez-Cardus A, et al: Epigenomic analysis detects aberrant super-enhancer DNA methylation in human cancer. Genome Biol 2016, 17:11.

