## Additional file 1 for "MethylationToActivity: a deep-learning framework that reveals promoter activity landscapes from DNA methylomes in individual tumors"

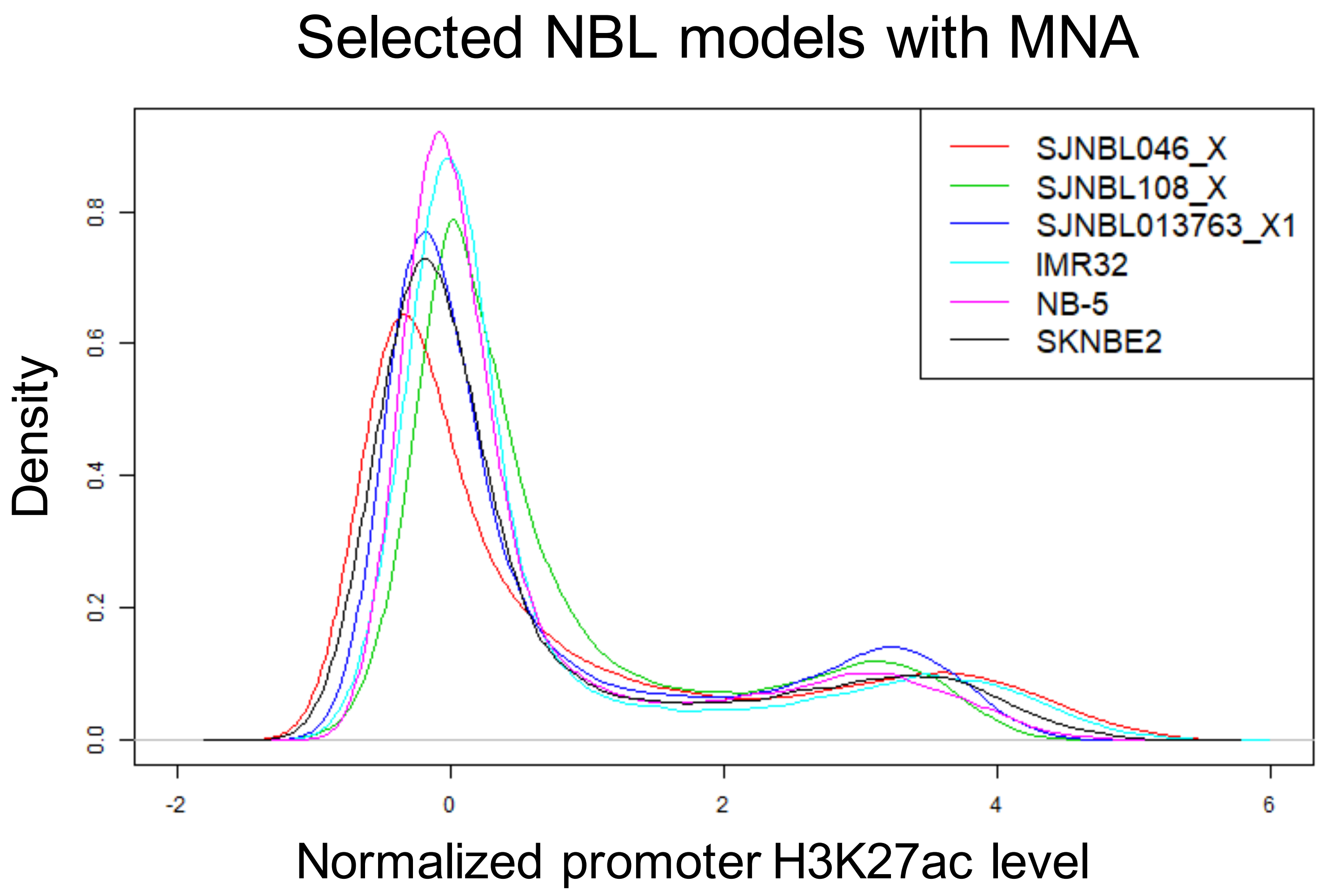

Selected NBL models with MNA
Density
0.0
0.2
0.4
0.6
0.8
-2
0
2
4
6
Normalized promoter H3K27ac level
SJNBL046_X
SJNBL108_X
SJNBL013763_X1
IMR32
NB-5
SKNBE2

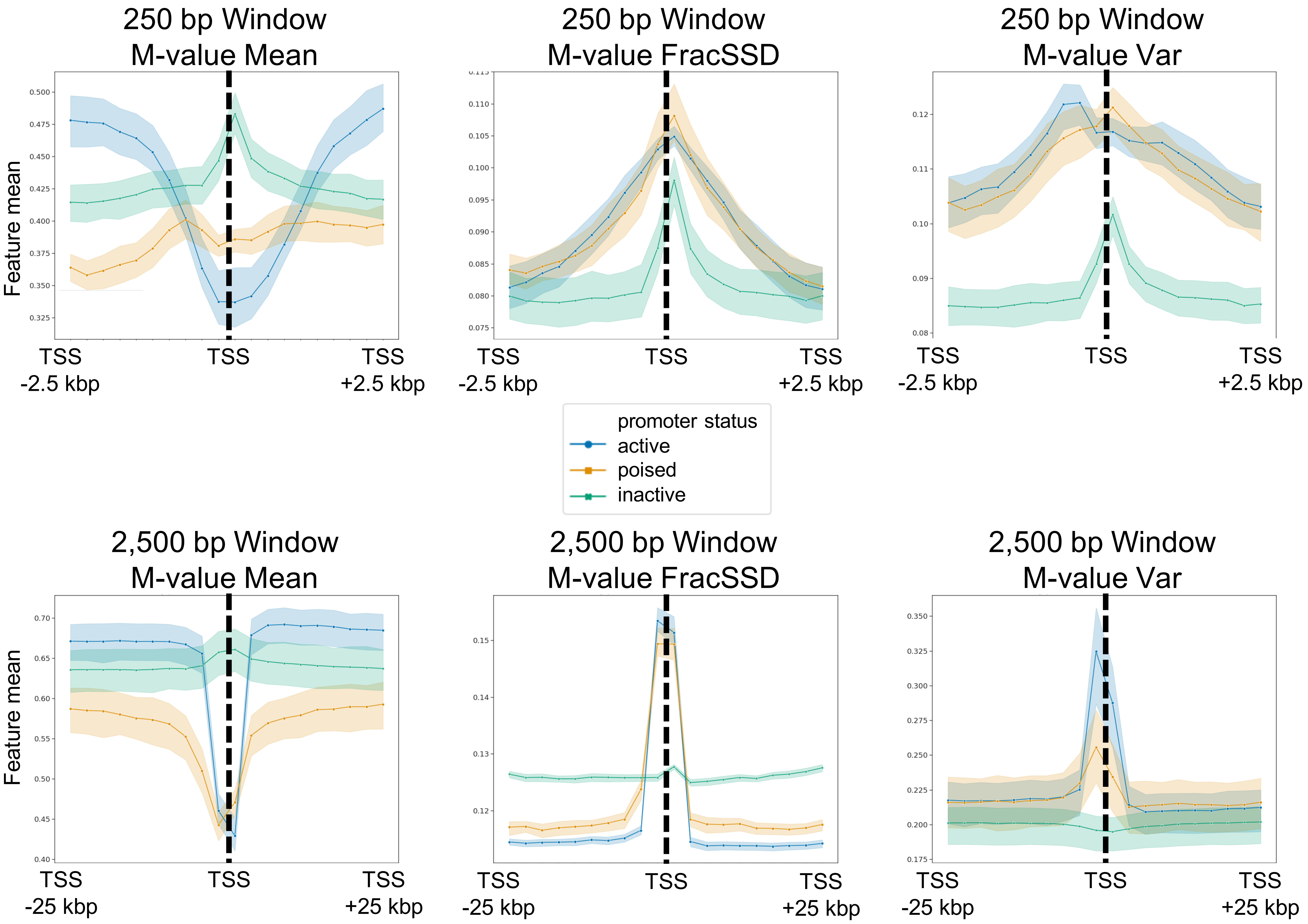
250 bp Window
M-value Mean
250 bp Window
M-value FracSSD
250 bp Window
M-value Var
Feature mean
TSS
-2.5 kbp
TSS
TSS
+2.5 kbp
promoter status
active
poised
inactive
2,500 bp Window
M-value Mean
2,500 bp Window
M-value FracSSD
2,500 bp Window
M-value Var
TSS
-25 kbp
TSS
TSS
+25 kbp

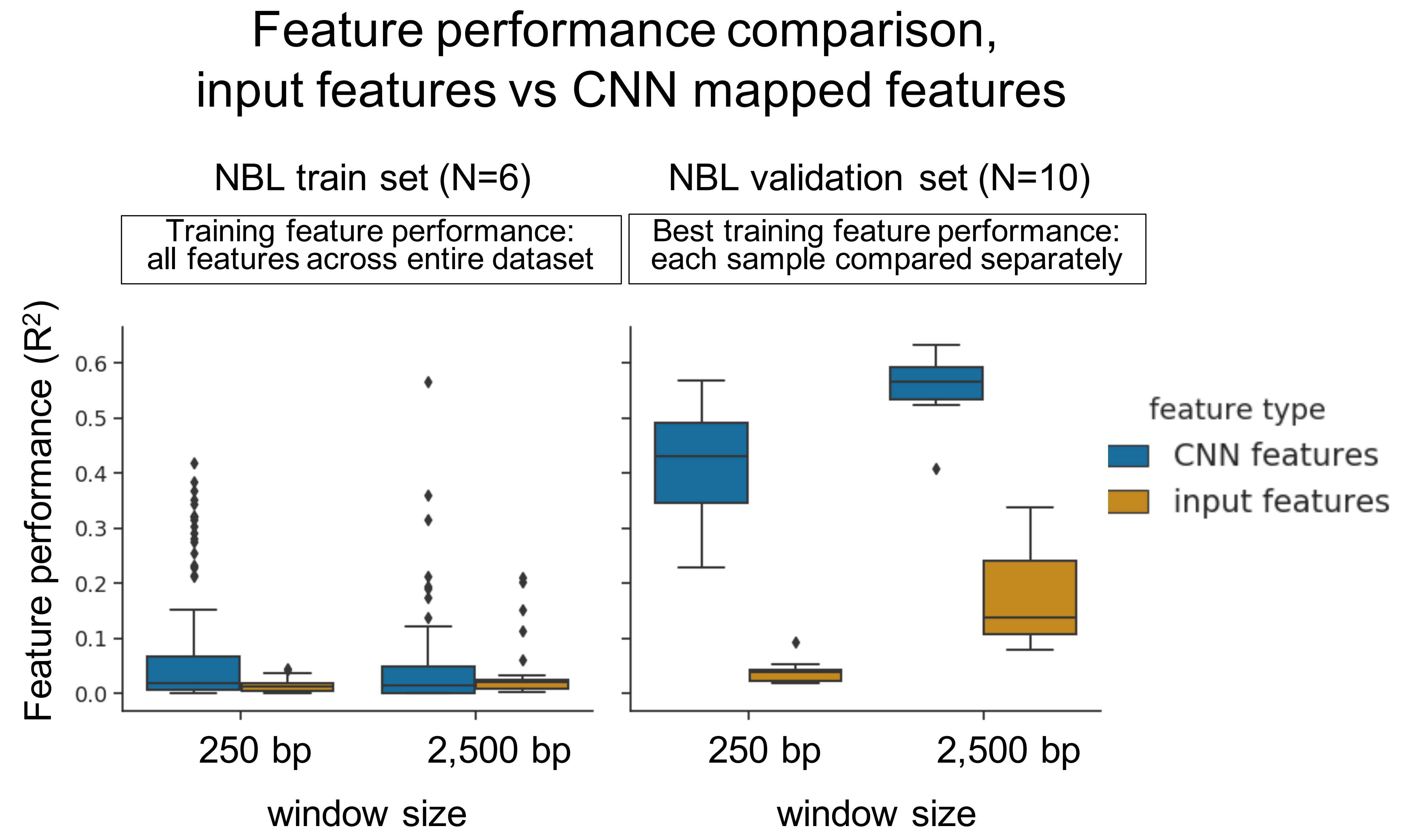
Feature performance comparison,
input features vs CNN mapped features
NBL train set (N=6)
NBL validation set (N=10)
Training feature performance:
all features across entire dataset
Best training feature performance:
each sample compared separately
Feature performance ($R^2$)
0.0
0.1
0.2
0.3
0.4
0.5
0.6
250 bp
2,500 bp
window size
feature type
CNN features
input features

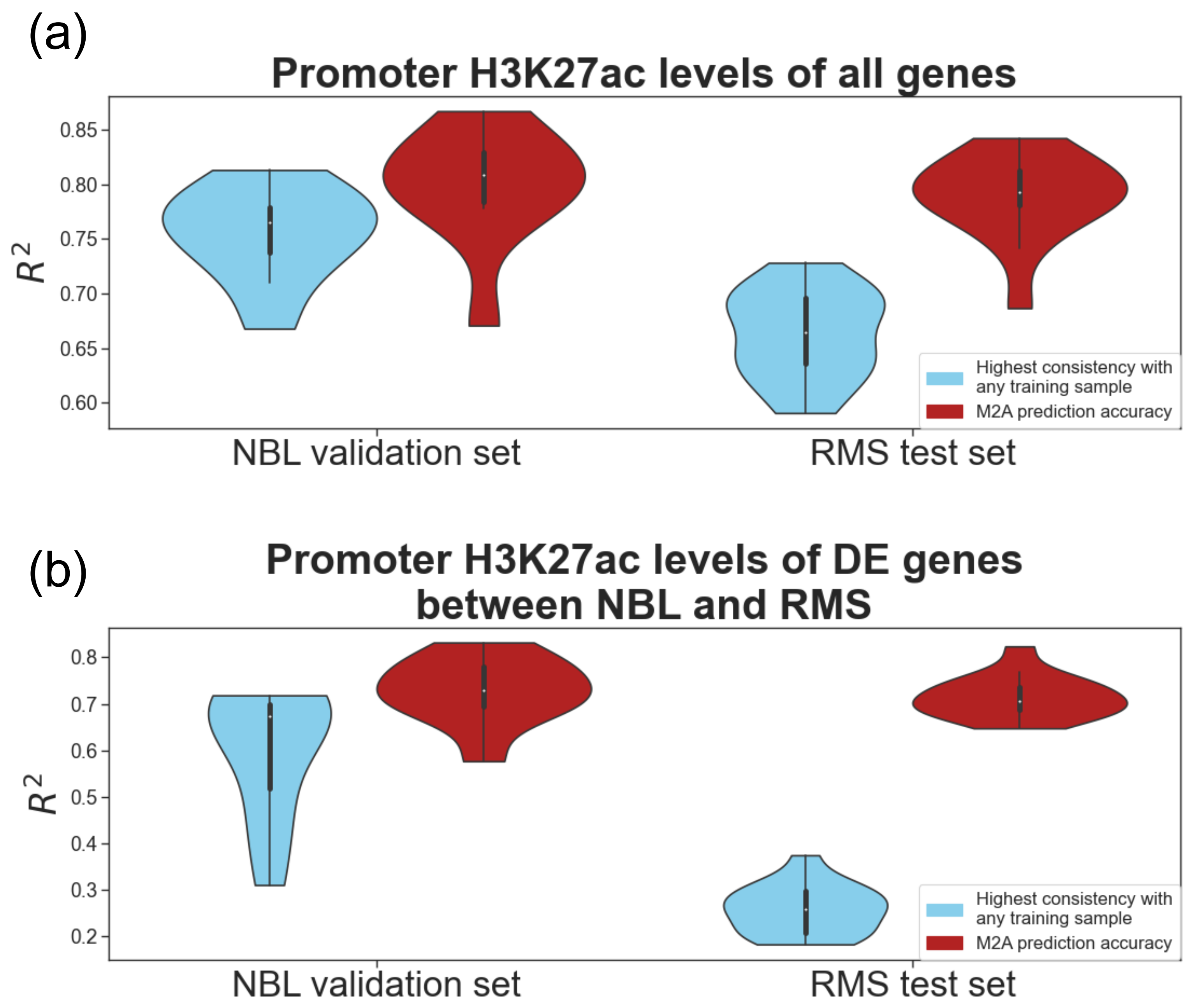
(a)
Promoter H3K27ac levels of all genes
$R^2$
0.85
0.80
0.75
0.70
0.65
0.60
Highest consistency with any training sample
M2A prediction accuracy
NBL validation set
RMS test set
(b)
Promoter H3K27ac levels of DE genes between NBL and RMS
$R^2$
0.8
0.7
0.6
0.5
0.4
0.3
0.2
Highest consistency with any training sample
M2A prediction accuracy
NBL validation set
RMS test set

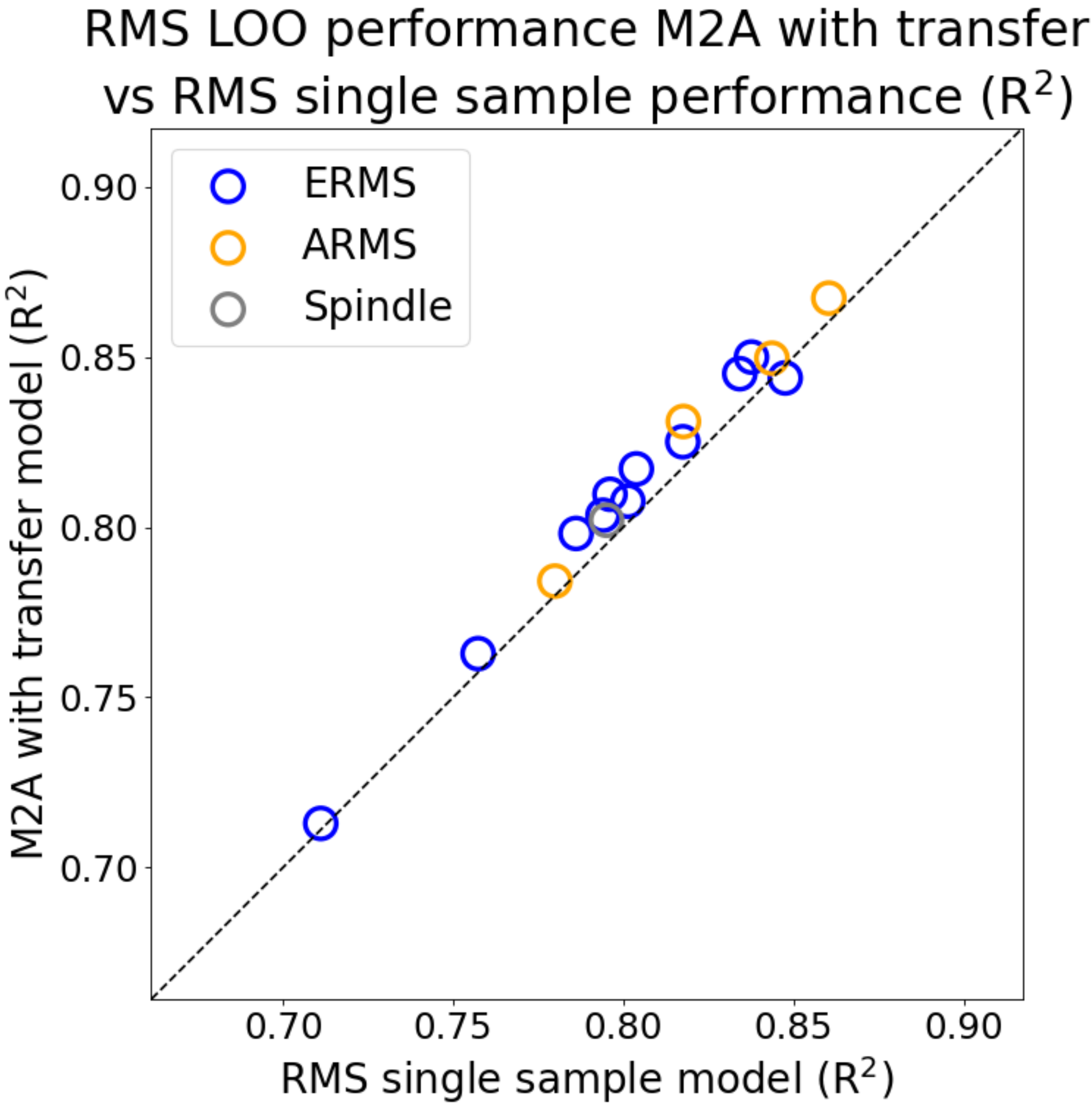

RMS LOO performance M2A with transfer vs RMS single sample performance ($R^2$)
ERMS
ARMS
Spindle
M2A with transfer model ($R^2$)
RMS single sample model ($R^2$)
0.70
0.75
0.80
0.85
0.90

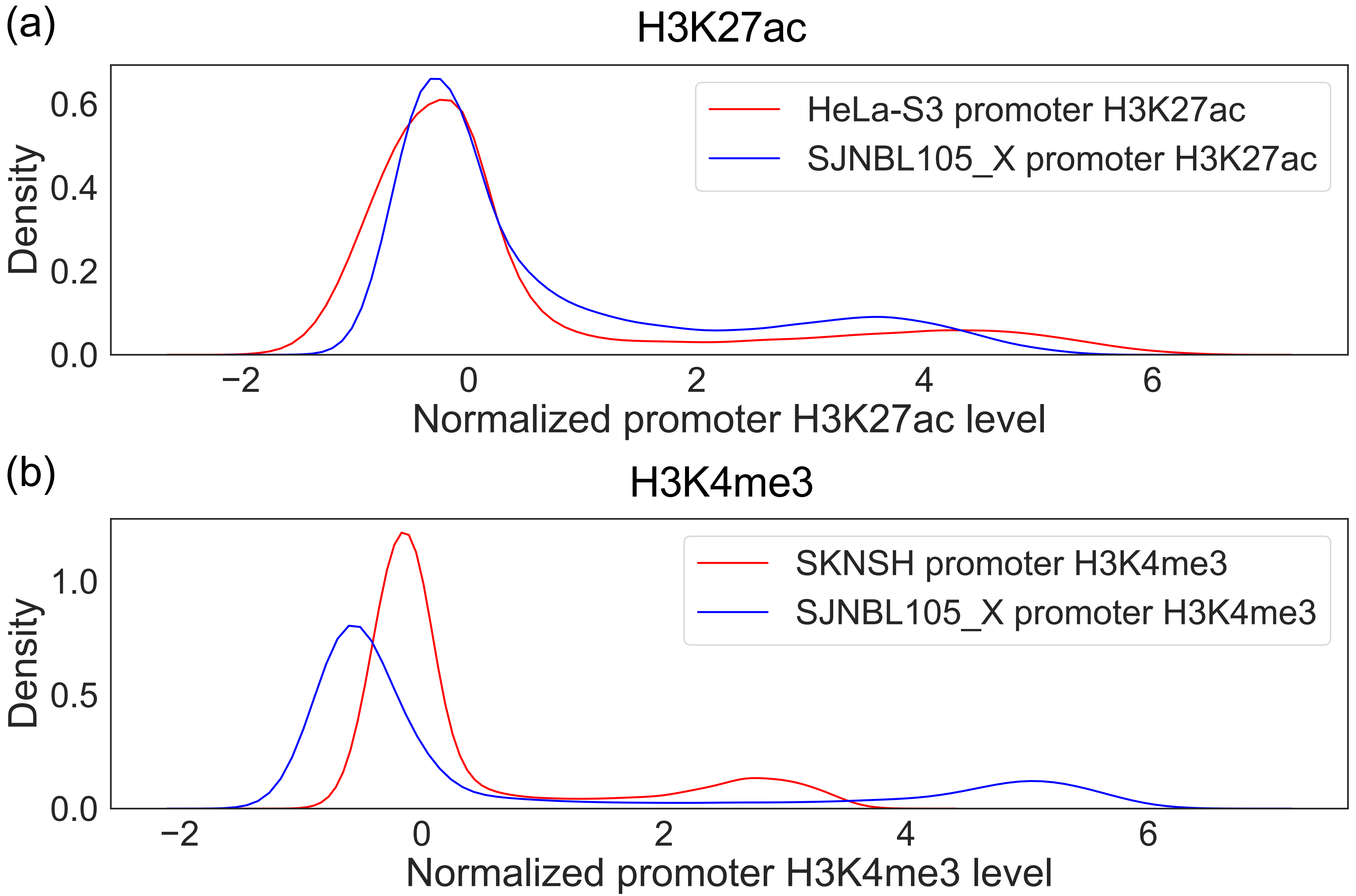

(a)
H3K27ac
HeLa-S3 promoter H3K27ac
SJNBL105_X promoter H3K27ac
Density
0.0
0.2
0.4
0.6
−2
0
2
4
6
Normalized promoter H3K27ac level
(b)
H3K4me3
SKNSH promoter H3K4me3
SJNBL105_X promoter H3K4me3
Density
0.0
0.5
1.0
−2
0
2
4
6
Normalized promoter H3K4me3 level

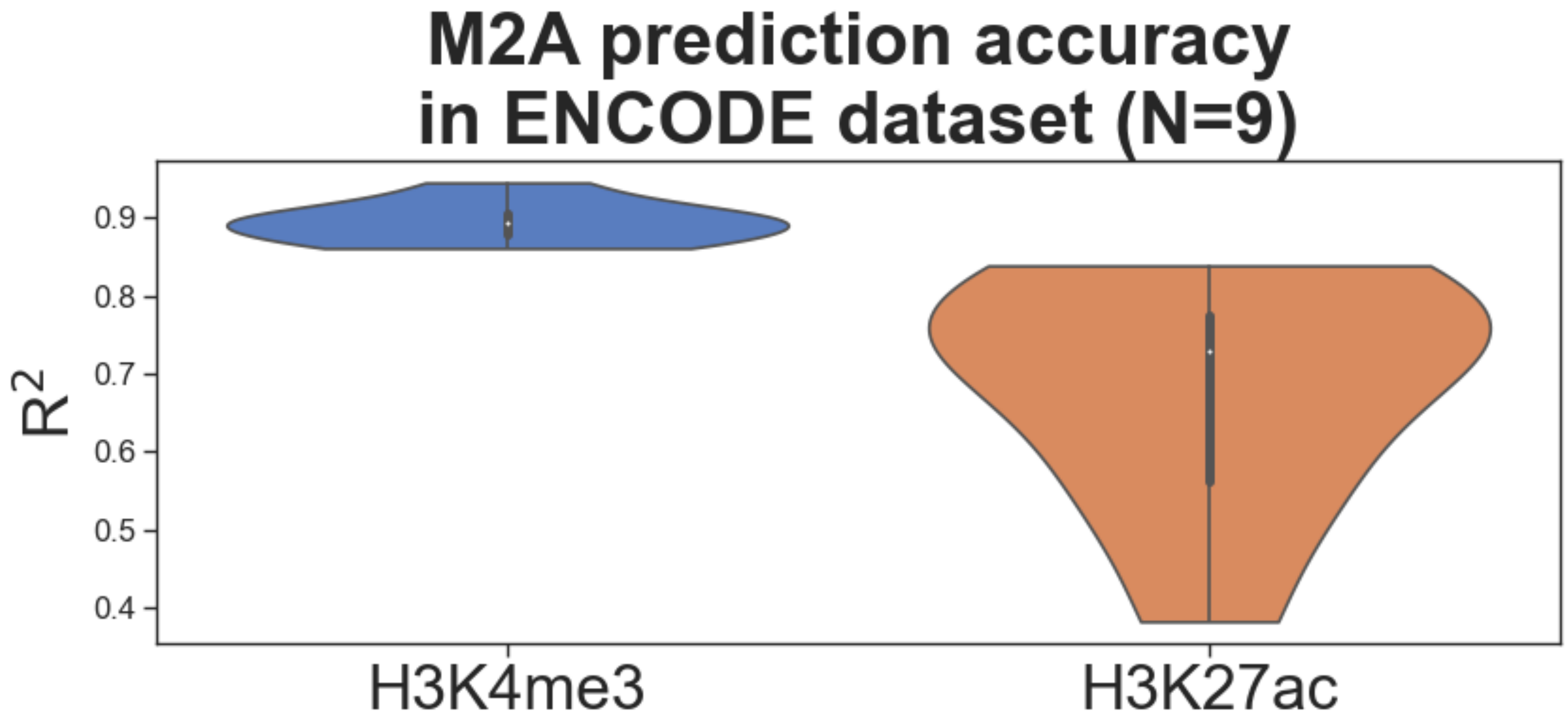

M2A prediction accuracy
in ENCODE dataset (N=9)
0.9
0.8
0.7
0.6
0.5
0.4
R2
H3K4me3
H3K27ac

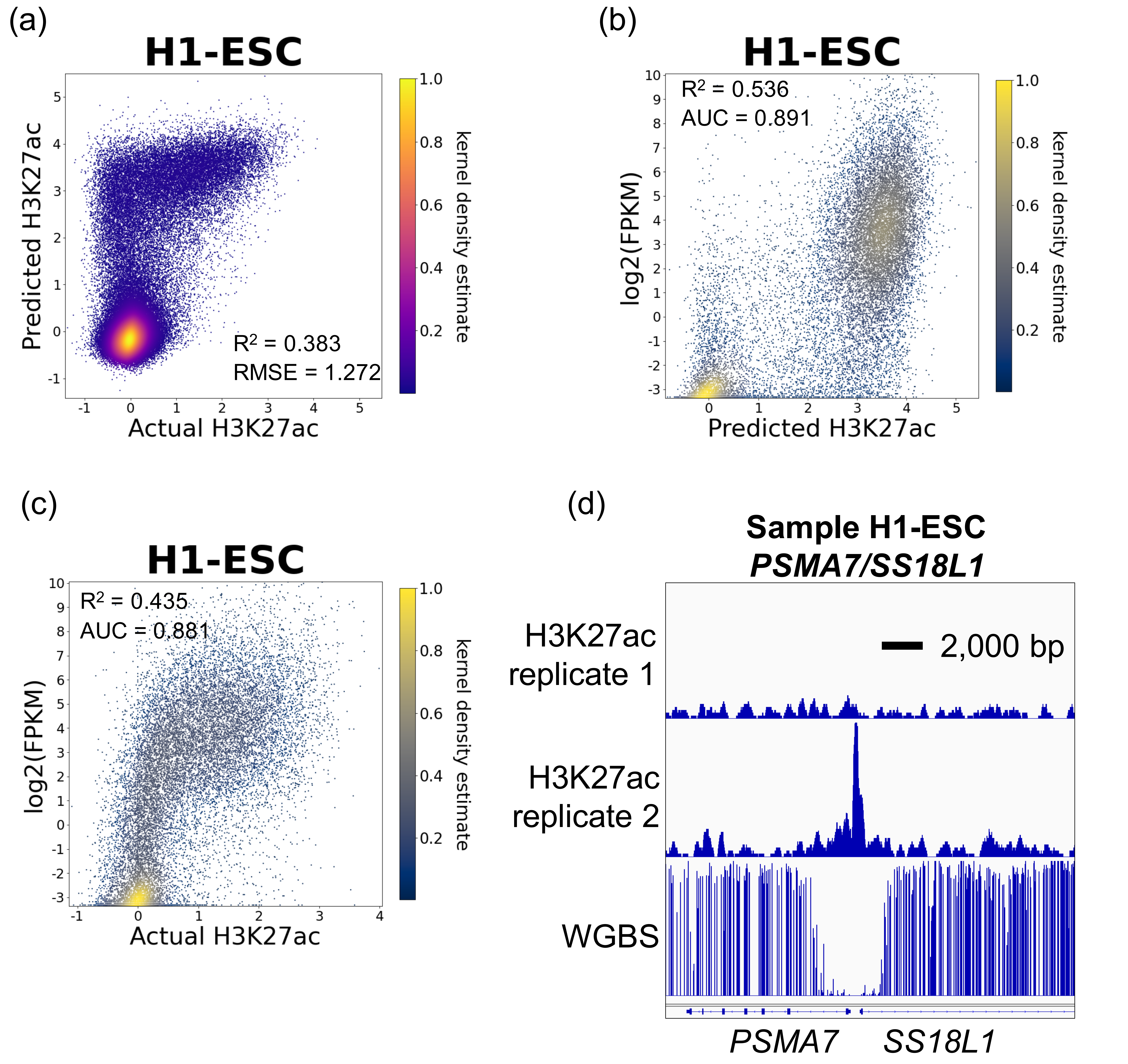

(a)
H1-ESC
Predicted H3K27ac
Actual H3K27ac
$R^2$ = 0.383
RMSE = 1.272
kernel density estimate
(b)
H1-ESC
log2(FPKM)
Predicted H3K27ac
$R^2$ = 0.536
AUC = 0.891
kernel density estimate
(c)
H1-ESC
log2(FPKM)
Actual H3K27ac
$R^2$ = 0.435
AUC = 0.881
kernel density estimate
(d)
Sample H1-ESC
*PSMA7/SS18L1*
H3K27ac replicate 1
2,000 bp
H3K27ac replicate 2
WGBS
*PSMA7*
*SS18L1*

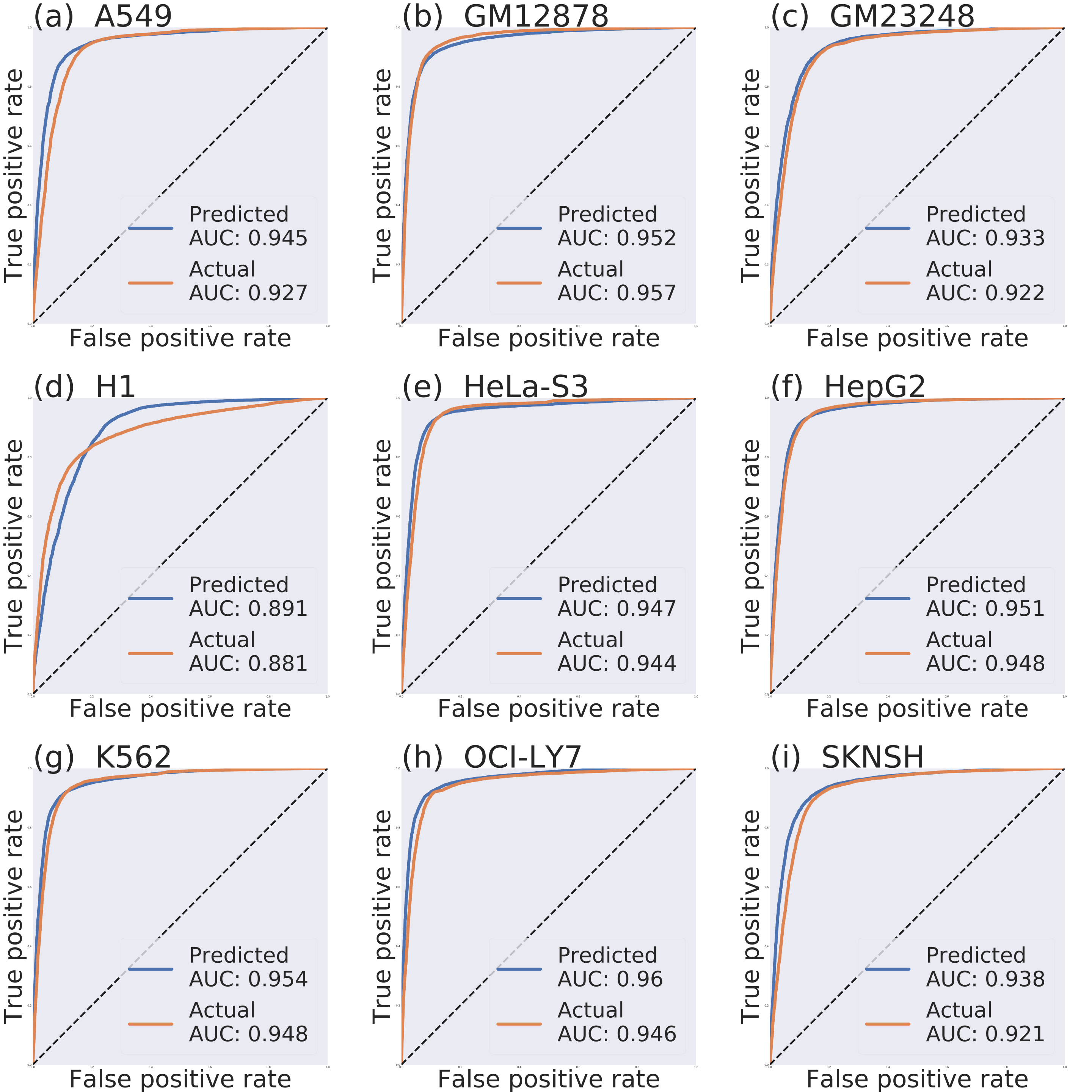
(a) A549
Predicted AUC: 0.945
Actual AUC: 0.927
(b) GM12878
Predicted AUC: 0.952
Actual AUC: 0.957
(c) GM23248
Predicted AUC: 0.933
Actual AUC: 0.922
(d) H1
Predicted AUC: 0.891
Actual AUC: 0.881
(e) HeLa-S3
Predicted AUC: 0.947
Actual AUC: 0.944
(f) HepG2
Predicted AUC: 0.951
Actual AUC: 0.948
(g) K562
Predicted AUC: 0.954
Actual AUC: 0.948
(h) OCI-LY7
Predicted AUC: 0.96
Actual AUC: 0.946
(i) SKNSH
Predicted AUC: 0.938
Actual AUC: 0.921
True positive rate
False positive rate

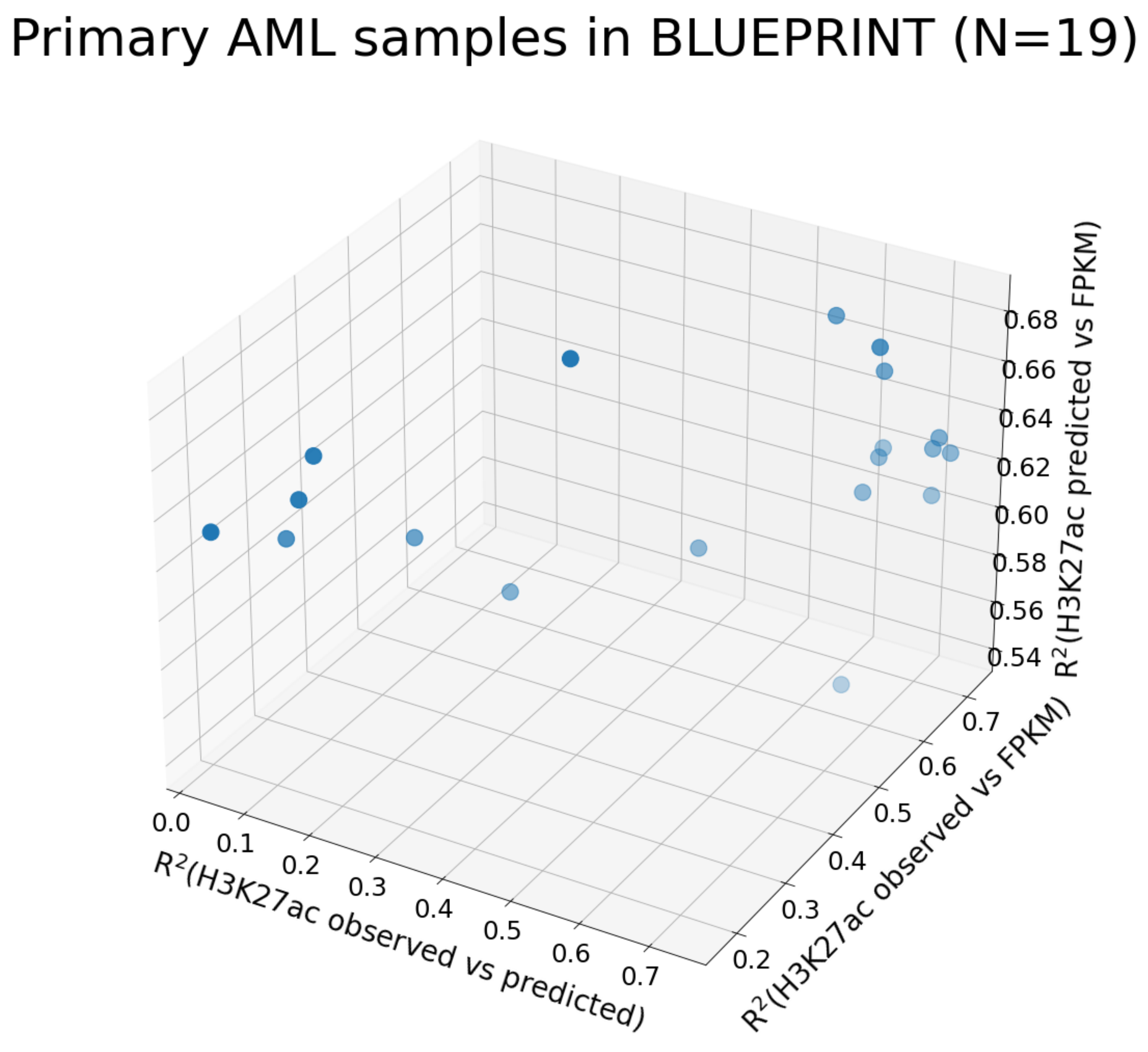

Primary AML samples in BLUEPRINT (N=19)
0.68
0.66
0.64
0.62
0.60
0.58
0.56
0.54
R²(H3K27ac predicted vs FPKM)
0.0
0.1
0.2
0.3
0.4
0.5
0.6
0.7
R²(H3K27ac observed vs predicted)
0.2
0.3
0.4
0.5
0.6
0.7
R²(H3K27ac observed vs FPKM)

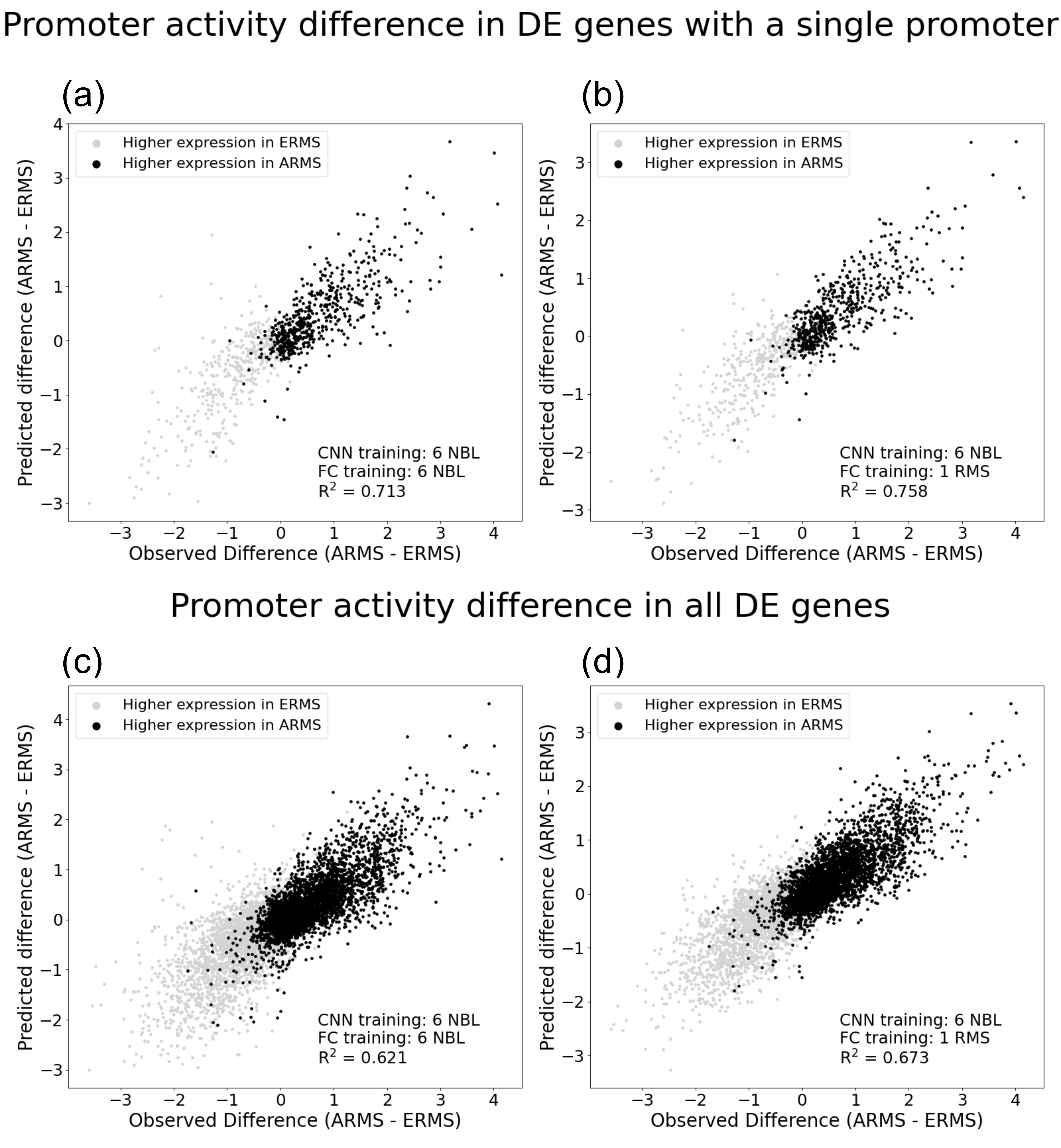
Promoter activity difference in DE genes with a single promoter
(a)
Higher expression in ERMS
Higher expression in ARMS
CNN training: 6 NBL
FC training: 6 NBL
R2 = 0.713
Predicted difference (ARMS - ERMS)
Observed Difference (ARMS - ERMS)
(b)
Higher expression in ERMS
Higher expression in ARMS
CNN training: 6 NBL
FC training: 1 RMS
R2 = 0.758
Predicted difference (ARMS - ERMS)
Observed Difference (ARMS - ERMS)
Promoter activity difference in all DE genes
(c)
Higher expression in ERMS
Higher expression in ARMS
CNN training: 6 NBL
FC training: 6 NBL
R2 = 0.621
Predicted difference (ARMS - ERMS)
Observed Difference (ARMS - ERMS)
(d)
Higher expression in ERMS
Higher expression in ARMS
CNN training: 6 NBL
FC training: 1 RMS
R2 = 0.673
Predicted difference (ARMS - ERMS)
Observed Difference (ARMS - ERMS)

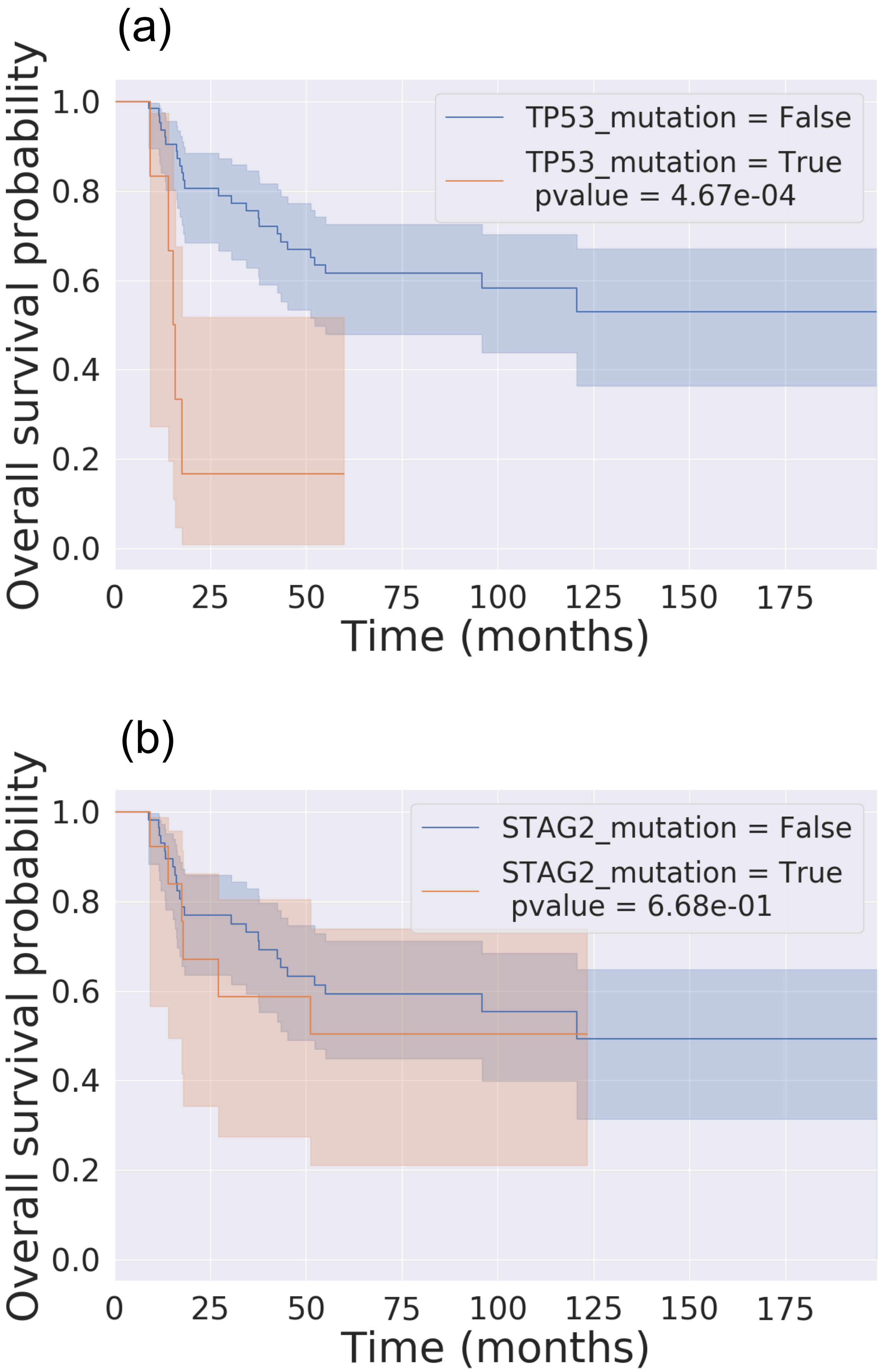

(a)
Overall survival probability
1.0
0.8
0.6
0.4
0.2
0.0
0
25
50
75
100
125
150
175
Time (months)
TP53_mutation = False
TP53_mutation = True
pvalue = 4.67e-04
(b)
Overall survival probability
1.0
0.8
0.6
0.4
0.2
0.0
0
25
50
75
100
125
150
175
Time (months)
STAG2_mutation = False
STAG2_mutation = True
pvalue = 6.68e-01

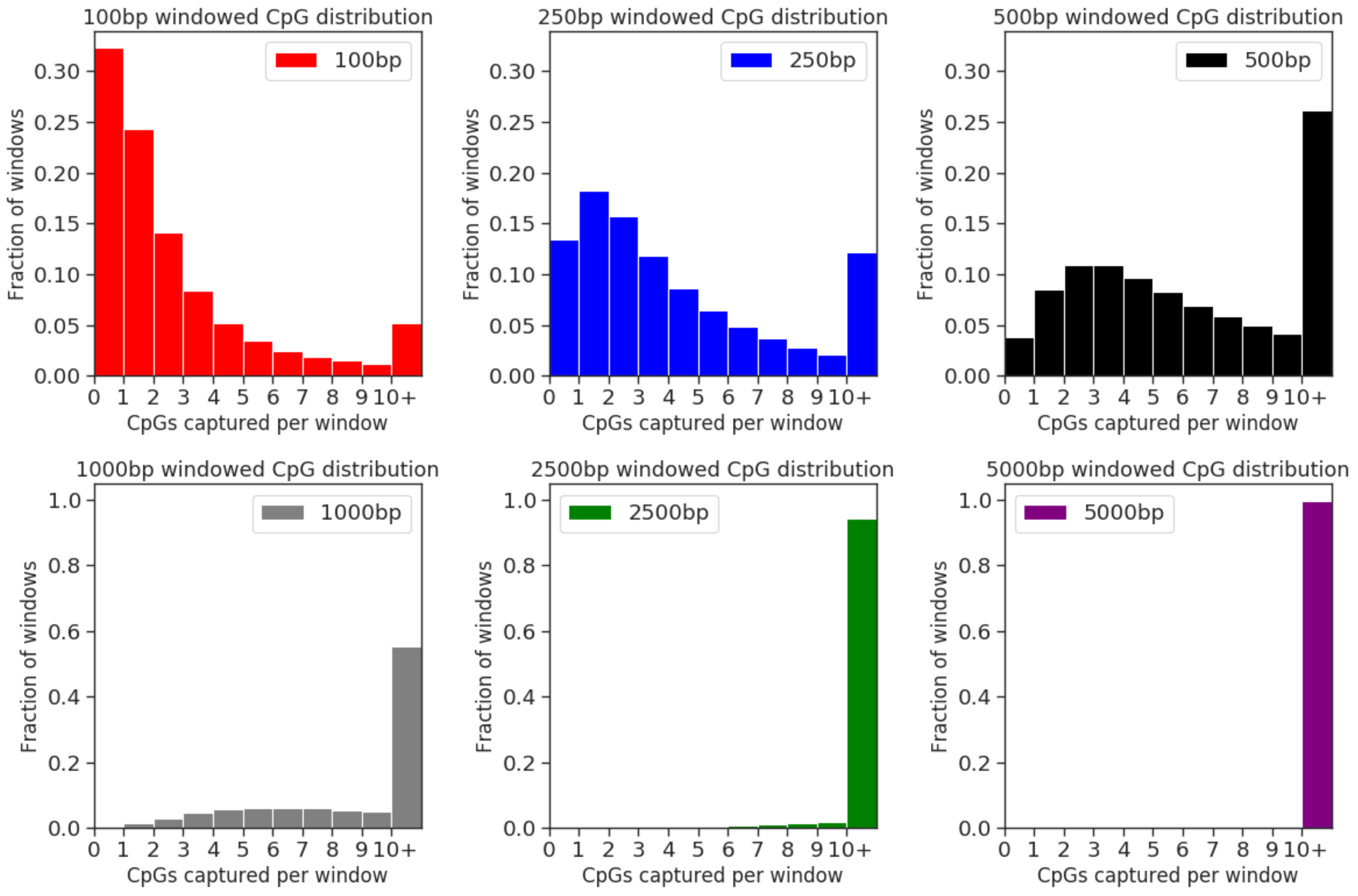
100bp windowed CpG distribution
100bp
250bp windowed CpG distribution
250bp
500bp windowed CpG distribution
500bp
1000bp windowed CpG distribution
1000bp
2500bp windowed CpG distribution
2500bp
5000bp windowed CpG distribution
5000bp
Fraction of windows
CpGs captured per window
0 1 2 3 4 5 6 7 8 9 10+
